# Temporal dynamics of base excision / single-strand break repair protein complex assembly and disassembly are modulated by the PARP1/NAD^+^/SIRT6 axis

**DOI:** 10.1101/2021.04.01.437913

**Authors:** Christopher A. Koczor, Kate M. Saville, Joel F. Andrews, Jennifer Clark, Qingming Fang, Jianfeng Li, Mikhail V. Makarov, Marie Migaud, Robert W. Sobol

## Abstract

Assembly and disassembly of DNA repair protein complexes at sites of DNA damage is essential to maintain genomic integrity. We investigated factors coordinating assembly of the base excision repair (BER) proteins, DNA polymerase β (Polβ) and XRCC1, to DNA lesion sites, identifying a new role for Polβ in regulating XRCC1 disassembly from DNA repair complexes and conversely, demonstrating Polβ’s dependence on XRCC1 for complex assembly. RealPAR, a genetically-encoded probe for live cell imaging of poly(ADP-ribose) (PAR), reveals that Polβ and XRCC1 require PAR for repair complex assembly and PAR degradation for disassembly. We find that BER complex assembly is further modulated by attenuation / augmentation of NAD^+^ biosynthesis. Finally, SIRT6 does not regulate PARP1 activation but impairs XRCC1 recruitment, leading to diminished Polβ abundance at sites of DNA damage. These findings highlight coordinated yet independent roles for both PARP1 and SIRT6 and their regulation by NAD^+^ bioavailability to facilitate BER.

## INTRODUCTION

DNA damage-induced genomic instability is a well-documented phenotype in both aging and cancer. Multiple DNA repair pathways facilitate repair of the damage, with each pathway specializing in the repair of specific DNA lesions. These repair pathways rely on the coordinated expression, synthesis and post-translational modification of multiple proteins and the bioavailability of regulatory factors to repair the DNA lesion, including 1) signaling to promote chromatin access and DNA repair complex assembly, 2) localization of repair complex scaffold proteins, 3) activity of enzymatic repair proteins, 4) disassembly of the repair complex and finally, 5) chromatin reorganization.

Base excision repair (BER) and single-strand break repair (SSBR) mechanisms facilitate repair of base damage and DNA single-strand breaks (Abbotts and Wilson, 2017; Svilar et al., 2011) (**Figure 1A**). Short-patch BER begins with removal of the damaged base by a DNA glycosylase to hydrolyze the N-glycosidic bond between the nucleobase and the phosphodiester riboside backbone. The resulting apurinic/apyrimidinic (AP) site is processed 5’ to the AP site by AP-endonuclease 1 (APE1) and the resulting gap is tailored by the 5’-deoxyribosephosphate (dRP) lyase function of DNA polymerase β (Polβ) (Sobol et al., 2000; Wilson and Barsky, 2001). This yields a single-strand break that is processed by Polβ’s nucleotidyl-transferase activity to insert a new base, followed by DNA ligase 1 (LIG1) or DNA ligase 3 (LIG3)-mediated ligation to seal the phosphodiester backbone. In the case of SSBR, a single strand break (similar to the intermediate observed during BER) undergoes gap tailoring by APE1 and Polβ to ensure the break is a substrate for new base addition by Polβ. Short-patch BER and SSBR can diverge depending on the 5’ and 3’ terminal blocking lesions and whether they are substrates for Polβ and APE1 (Abbotts and Wilson, 2017).

**Figure 1:**
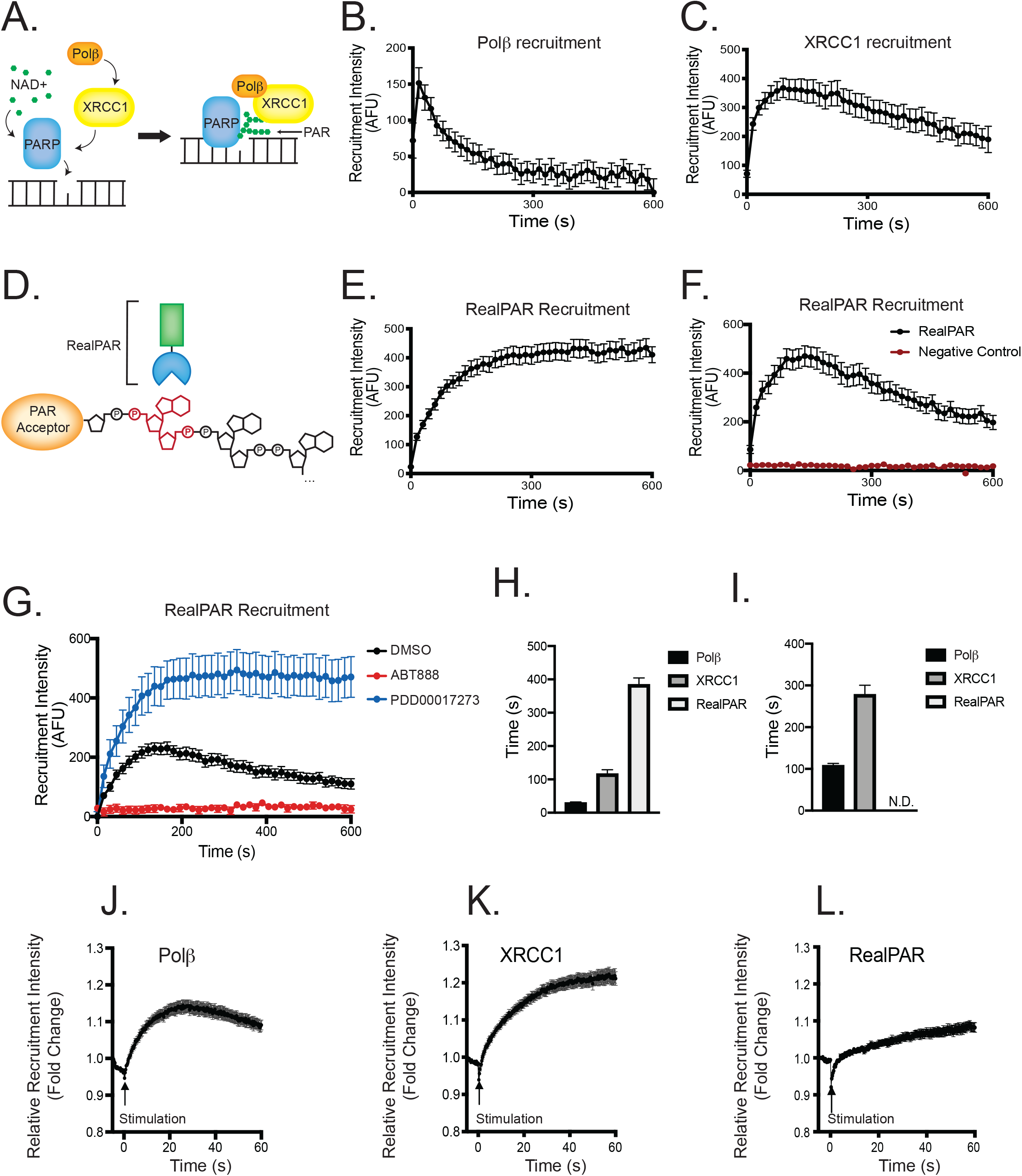
Laser-induced micro-irradiation (UVA-355nm) of Polβ, XRCC1 and RealPAR. A) Current model for Polβ/XRCC1/PAR complex formation. B) Recruitment of EGFP-Polβ to sites of laser micro-irradiation (355nm) in U2OS cells, n ≥35. Error bars indicate standard error of the mean. C) Recruitment of XRCC1-EGFP to sites of laser micro-irradiation (355nm) in U2OS cells, n ≥35. D) Model of RealPAR’s mode of action. RealPAR contains a poly(ADP-ribose) (PAR) binding motif fused to EGFP. Upon micro-irradiation, RealPAR binds to the iso-ADP-ribose moiety (shown in red) of PAR formed at sites of DNA damage. E) Recruitment of RealPAR to sites of laser micro-irradiation (355nm) in U2OS cells, n ≥35. F) Recruitment of RealPAR to sites of laser micro-irradiation (355nm) in A549 cells, n ≥35. A single point mutant (Y107A) in RealPAR’s PAR binding motif prevented focal recruitment. G) Inhibition of PARP or PARG alters RealPAR recruitment to sites of laser micro-irradiation in A549 cells, n ≥35. PARP inhibition, using velipirab (ABT-888, 10μM), prevented RealPAR’s recruitment, while PARG inhibition (PDD00017273, 10μM) retained RealPAR at sites of micro-irradiation by inhibiting the PARG-mediated degradation of PAR. H) Time to peak recruitment intensity of Polβ, XRCC1, and RealPAR in U2OS cells, n ≥35. Polβ reached peak recruitment at 30 ± 2s (mean ± SEM), XRCC1 at 116 ± 13s, and RealPAR at 384 ± 20s. Error bars indicate standard error of the mean. I) Half-life of recruitment of Polβ, XRCC1, and RealPAR in U2OS cells, n ≥35. The half-life of recruitment was 108 ± 5s (mean ± SEM) for Polβ, 278 ± 22s for XRCC1, and was not detected for RealPAR for 20 minutes post-stimulation in U2OS cells. J) Serial micro-irradiation of EGFP-Polβ in U2OS cells, n ≥40. A 250μs imaging interval was used to image single cells with higher temporal resolution than in parallel imaging. Results confirm no recruitment peak was observed earlier than the peak observed in the parallel mode results in Figure 1B. Error bars indicate standard error of the mean. K) Serial micro-irradiation of XRCC1-EGFP in U2OS cells, n ≥40. L) Serial micro-irradiation of RealPAR in U2OS cells, n ≥40. See also Figure S1.

For the efficient recruitment of BER/SSBR proteins to sites of damage, multiple factors must work in coordination. X-ray repair cross complementing 1 (XRCC1) functions as a scaffold protein that localizes repair proteins (including Polβ and DNA ligase 3) to DNA damage sites, thereby facilitating BER/SSBR repair (Almeida and Sobol, 2007; London, 2015). XRCC1 is recruited through its poly(ADP-ribose) (PAR) binding domain, which binds to PAR chains formed by activated poly(ADP-ribose) polymerases (PARPs) at sites of DNA damage (Breslin et al., 2015; El-Khamisy et al., 2003). Loss of one or more of these assembly intermediates or proteins would be expected to compromise BER/SSBR complex formation by reducing Polβ localization to sites of DNA damage. This appears to be the case for NAD^+^-dependent protein deacetylase sirtuin-6 (SIRT6). Loss of SIRT6 in mammalian cells and mice induces increased genomic instability and enhanced sensitivity to DNA alkylation and oxidation damage (Mostoslavsky et al., 2006). The mechanism for this instability has not been fully characterized, although it has been shown that enhancing Polβ’s dRP lyase function through over-expression rescued cellular sensitivity to DNA alkylation damage (Mostoslavsky et al., 2006).

Another factor critical to genome stability is the co-factor nicotinamide adenine dinucleotide (NAD^+^) (Fouquerel and Sobol, 2014). A deficiency in NAD^+^ can lead to decreased PARP and sirtuin activity, promoting genomic instability and a decrease in DNA repair capacity (Fouquerel et al., 2014). The cofactor NAD^+^ is a required substrate to enable PARP-mediated PAR formation at sites of DNA damage (Rouleau et al., 2010). Additionally, NAD^+^ is required by sirtuins (such as SIRT6) to perform a number of genome stabilizing activities (Imai and Guarente, 2014). NAD^+^ levels have been shown to deplete with aging (Fang et al., 2017; Imai and Guarente, 2014), during pregnancy (Shi et al., 2017), upon viral infection (Mesquita et al., 2016), and are dysregulated in some cancers (Chiarugi et al., 2012; Yaku et al., 2018). Augmenting NAD^+^ through pharmacological means has been of increasing interest (Giroud-Gerbetant et al., 2019). Dihydronicotinamide riboside (NRH) is a reduced form of nicotinamide riboside (NR) and is uniquely metabolized, leading to enhanced levels of intracellular NAD^+^ (Giroud-Gerbetant et al., 2019; Yang et al., 2019). Currently, it is unknown how modulating NAD^+^ impacts assembly and disassembly of BER/SSBR complexes at sites of DNA damage.

While the *in vitro* biochemistry of the proteins in the BER and SSBR pathways have been extensively studied, the mechanisms by which key repair proteins assemble and disassemble at the DNA damage site in the cell are not defined. Biochemical analysis of BER processing of DNA lesions using purified nucleosomes suggests additional factors are likely required to effectively gain access to the DNA lesion for removal and repair (Beard et al., 2003; Cole et al., 2010; Rodriguez et al., 2017). Laser micro-irradiation provides real-time assessment of DNA repair in live cells using fluorescently tagged fusion proteins within the cellular context of other factors known to alter DNA repair, including chromatin structure, non-enzymatic accessory proteins, and the cellular metabolic profile. Here, real-time *in vivo* assembly and disassembly of BER/SSBR complexes were investigated using UVA (355nm) laser micro-irradiation to introduce lesions repaired primarily through BER/SSBR (Holton et al., 2017). Knockout of critical proteins demonstrates key steps for Polβ and XRCC1 recruitment to and release from sites of micro-irradiation-induced DNA damage. To follow PARP1-activation in real-time, we developed the RealPAR probe, a genetically-encoded PAR monitor, demonstrating an enhanced capability to extend the characterization of BER/SSBR to include real-time PAR formation *in vivo* (in cells) and its effects on Polβ and XRCC1 recruitment. Further, we show that BER/SSBR complex assembly modulated by alterations in NAD^+^ bioavailability. Finally, we highlight that, unlike its role in DNA double-strand break repair (DSBR) (Tian et al., 2019), SIRT6 does not regulate PARP1 activation during BER yet negatively impacts XRCC1’s complex assembly capacity (independent of PAR formation) and ultimately reduces Polβ’s localization to sites of DNA damage. Overall, these studies highlight the coordinated yet independent roles for both PARP1 and SIRT6 and their regulation by NAD^+^ to facilitate BER, supporting an essential PARP1-NAD^+^-SIRT6 axis for BER protein complex assembly dynamics.

## RESULTS

### Dynamics of Polβ and XRCC1 base excision repair complex assembly

To quantitatively assess the recruitment of DNA repair proteins in response to laser-induced micro-irradiation, we used **MIDAS** (**M**odular **I**rradiation, **D**etection, and **A**nalysis **S**ystem), a software platform for performing and analyzing micro-irradiation experiments (see Methods). To rapidly identify the recruitment kinetic profiles for individual fluorescently tagged proteins, we utilized the parallel stimulation module in MIDAS to collect time-lapse images of multiple laser-induced DNA damage foci (one per cell nucleus) for quantitative and statistical analysis, irradiating 10 cells per field and acquiring an image every 15 seconds for the duration of the study (either 10 or 20 minutes). Although cells in each image field are irradiated sequentially, cell-specific timing offsets are measured for each irradiation event, allowing for precise calculation of timing on a per-cell basis.

To investigate the Polβ/XRCC1/PAR axis of base excision repair (BER) (**Figure 1A**), we established recruitment kinetics for the central protein factors in BER, Polβ and XRCC1. Previous studies have demonstrated that 405nm laser micro-irradiation introduces both DNA single-strand breaks (SSB) and DNA double-strand breaks (DSB), while 355nm lasers can produce BER and SSBR-specific damage (Holton et al., 2017; Lan et al., 2004). Cells expressing the fluorescently tagged DNA double-strand break repair (DSBR) protein 53BP1 are readily recruited to sites of DNA damage induced by 405nm micro-irradiation. However, 53BP1 did not respond to 355nm micro-irradiation, verifying at those settings that the 355nm laser does not produce measurable recruitment of DSB-responsive proteins (**Figures S1A and S1B**). Thus, using 355nm stimulation allows investigation of the response of Polβ and XRCC1 to BER/SSBR selective DNA damage in the absence of significant DNA double-stranded break repair and response from DSBR proteins.

Polβ and XRCC1 recruited rapidly to 355nm laser-induced DNA damage in U2OS cells (**Figure 1B, C, and S1C**). Previous research by our lab and others supports a paradigm where Polβ and XRCC1 form a heterodimer during DNA repair, which suggests similar DNA damage-induced recruitment profiles (Almeida and Sobol, 2007; Fang et al., 2019; Fang et al., 2014). However, Polβ was found to reach maximum recruitment capacity more rapidly (30s to peak intensity) than XRCC1 (∼116s). Recruitment of Polβ and XRCC1 was verified in a second cell line (A549 cells), though recruitment peaks were delayed for both repair proteins in the A549 cell line, demonstrating the importance of cell-type specific context for repair complex assembly and disassembly (**Figures S1D-E**).

### RealPAR enables real-time, live cell poly(ADP-ribose) (PAR) imaging

Due to the significance of PAR synthesis in promoting BER/SSBR complex assembly at sites of DNA damage, immunocytochemistry and PAR antibodies are routinely utilized to verify PAR formation at the site of damage, coincident with EGFP-Polβ recruitment (**Figure S1F**). However, this method has multiple drawbacks when compared to live-cell fluorescence imaging, including experimental variability in fixation and labeling methods as well as reduced temporal resolution due to cell fixation time. We developed a novel fluorescently tagged PAR-detecting fusion protein referred to as RealPAR (**Figure 1D**). Nine PAR binding domains (PBD) from known PAR-binding proteins were identified, with domains known to bind to different PAR chain moieties (**Supplemental Table S1**) (Teloni and Altmeyer, 2016). Each PBD was fused to a C-terminal EGFP domain, expressed in cells, and imaged to ensure expression of the fusion protein and validate sub-cellular localization, if any (**Figure S1G**). We visualized the response of each fusion protein to 355nm laser-induced DNA damage, under similar micro-irradiation conditions employed for Polβ and XRCC1 recruitment. Among the nine fusion proteins, only one (hereafter termed RealPAR) demonstrated recruitment to sites of 355nm micro-irradiation and was readily expressed in multiple cell types (**Figure 1E, F, and S1C**). Furthermore, 405nm micro-irradiation (which produces more DSBs but also elicits PAR formation) also promoted RealPAR’s recruitment to DNA damage-induced foci, verifying that RealPAR is recruited to at least two known PAR-generating micro-irradiation wavelengths (**Figure S1H**). The RealPAR PBD has been investigated previously, and mutations that eliminate PAR binding have been characterized (Wang et al., 2012). Expression of RealPAR harboring one of these mutations (Y107A) was sufficient to prevent RealPAR recruitment to micro-irradiation induced sites of DNA damage (**Figure 1F**). Finally, to determine if RealPAR’s recruitment was sensitive to and dependent on PAR formation and degradation, cells expressing RealPAR were treated with the PARP inhibitor ABT-888 or the PARG inhibitor PDD00017273 (Donawho et al., 2007; James et al., 2016). While PARP inhibition was able to prevent RealPAR’s recruitment to laser-induced foci, PARG inhibition enhanced and prolonged RealPAR’s recruitment (**Figure 1G**). RealPAR therefore is a stable, live-cell imaging tool for visualizing PAR formation and degradation in real-time in living cells. Combining RealPAR with laser micro-irradiation yields a powerful experimental platform for probing the mechanistic processes affecting DNA repair complex assembly (**Figure 1A**).

We used MIDAS to quantitatively assess recruitment, measuring the time to peak recruitment intensity and half-life of recruitment for Polβ, XRCC1, and RealPAR (**Figures 1H, I**). Polβ reached peak recruitment intensity first, followed by XRCC1 and then RealPAR. Similarly, the half-life of recruitment analysis demonstrated that Polβ disassembles from the repair complex first, followed by XRCC1 and the degradation of PAR, as evidenced by the long half-life of recruitment for RealPAR. To characterize the early phase of recruitment more precisely, we used MIDAS to perform high-speed single-cell micro-irradiation experiments, using an imaging interval of 250µs and a duration of 1 minute. This serial imaging approach was important for accurately characterizing Polβ, which exhibited the shortest time to peak intensity (**Figure 1B**). Serial analysis confirmed the 30s time to peak intensity for Polβ following 355nm stimulation in U2OS cells (**Figure 1J-L**).

### Over-expression of EGFP-Polβ displays similar recruitment kinetics as endogenously tagged EGFP-Polβ

While over-expression of fluorescently tagged DNA repair proteins can aid in understanding repair complex assembly kinetics, over-expression could alter repair kinetics by supplementing rate-limiting enzymes. This is important in the case of Polβ, which is required for short-patch BER, can be rate-limiting, and is the primary gap-tailoring dRP lyase for repair of base lesions (Allinson et al., 2001; Sobol et al., 2000; Srivastava et al., 1998). To address if over-expression altered recruitment kinetics, EGFP cDNA was fused endogenously to the POLB gene in chromosome 8 of A549 cells, thereby preserving the promoter region and allowing expression of the EGFP-Polβ protein under endogenous conditions (**Figure 2A**). We utilized a CRISPR/Cas9 ribonucleoprotein complex along with an oligonucleotide fragment of the POLB homology region around the POLB transcription start site and an in-frame EGFP cDNA. Successful generation of cells expressing EGFP-Polβ was confirmed through three independent methods. Sanger sequencing of POLB alleles demonstrated successful targeting of one out of the three alleles in A549 cells (**Figure 2B** and **Supplemental Table S2**). One allele had no modification, and the final allele had a partial incorporation that added 45 base pairs 5’ to POLB exon 1. Immunoblots showed that both EGFP-Polβ and non-tagged Polβ were produced by the modified A549 cells (**Figure 2C**; full blots in **Figure S2A**). Finally, we performed confocal spectral microscopy to confirm that EGFP-Polβ was detected in the modified A549 cells. Because expression under the endogenous promoter led to low levels of EGFP fluorescence in the modified cells, we performed spectral imaging and unmixing to remove autofluorescence. Spectral unmixing demonstrated that EGFP-Polβ expression was primarily in the nuclear compartment, with a minor fraction in the cytosolic compartment, consistent with the distribution observed in EGFP-Polβ overexpressing cells (**Figure 2D** **and S2B**). Finally, we performed 355nm micro-irradiation on the endogenously tagged A549/EGFP-Polβ cells. Endogenous EGFP-Polβ recruited to sites of 355nm laser-induced DNA damage, and recruitment kinetics were found to be similar to the recruitment kinetics of over-expressed EGFP-Polβ (**Figures 2D-G**). The half-life of recruitment for the endogenously tagged EGFP-Polβ was significantly reduced as compared to over-expressed EGFP-Polβ in A549 cells (**Figure 2G**). This may be a result of the increased amount of Polβ protein that can recruit to the site of damage in over-expression models, resulting in a slight increase in the time required to disassemble the Polβ complex. Overall, over-expression of EGFP-Polβ does not lead to gross changes in recruitment kinetics when compared to EGFP-Polβ expressed at endogenous levels.

**Figure 2:**
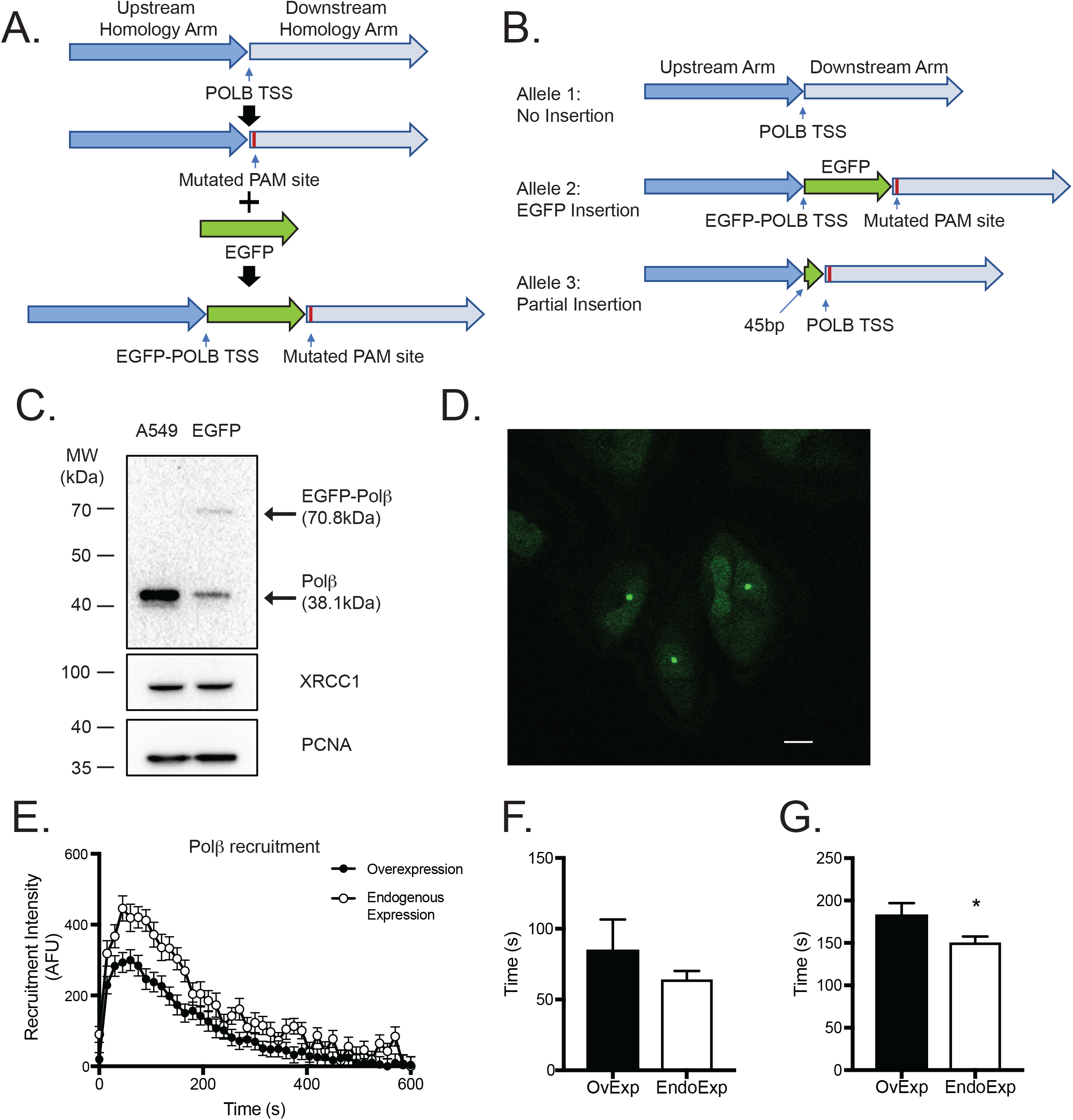
Over-expression of EGFP-Polβ displays similar recruitment kinetics as endogenously tagged EGFP-Polβ. A) Genomic editing strategy to target the POLB gene on chromosome 8 in A549 cells. The POLB homology region fragment (∼800bp on each side from the transcription start site) was amplified by high-fidelity PCR from A549 genomic DNA. EGFP cDNA was inserted in-frame with the transcriptional start site of POLB, and a silent mutation was placed at the PAM site (AGG to AAG) to prevent re-cleavage by CRISPR/Cas9. B) Allele sequencing results. Of the three alleles in A549 cells, one was not modified, one was modified with the full length EGFP in frame with POLB exon 1, and one allele displayed a partial 45bp insertion. The full sequencing results can be found in **Supplemental Table S2**. C) Immunoblot of endogenously tagged A549 cells. Bands were observed at 38.1 kDa (non-modified Polβ) and 70.8kDa (EGFP-Polβ) for the endogenous EGFP-Polβ cells. No changes in XRCC1 were observed. PCNA was used for even loading. D) Spectrally unmixed image of endogenously tagged EGFP-Polβ in A549 cells. Each cell displays both nuclear and cytosolic Polβ. The foci in the image demonstrates EGFP-Polβ recruitment to micro-irradiation induced DNA damage. E) Recruitment of endogenous EGFP-Polβ (open circles) and over-expressed EGFP-Polβ (closed circles) to sites of laser micro-irradiation (355nm), n ≥35. Error bars indicate standard error of the mean. F) Time to peak recruitment intensity of endogenous EGFP-Polβ and over-expressed EGFP-Polβ following laser micro-irradiation. No significant difference was observed following a Student’s t-test. Error bars indicate standard error of the mean. G) Half-life of recruitment of endogenous EGFP-Polβ and over-expressed EGFP-Polβ following laser micro-irradiation. A significant difference (p<0.05) was observed (Student’s t-test), with endogenous EGFP-Polβ disassembling from the repair complex faster than over-expressed EGFP-Polβ. See also Figure S2.

### Loss of Polβ enzymatic activity does not alter damage-induced recruitment kinetics

Polβ has two enzymatic functions: (i) a 5’dRP lyase activity that can be significantly attenuated by an alanine mutation at amino acid residue K72 (K72A) and (ii) a polymerase or nucleotidyl-transferase activity that can be eliminated by an alanine mutation at residue D256 (D256A) (Matsumoto et al., 1998; Menge et al., 1995). Loss of the 5’dRP lyase activity of Polβ, but not its polymerase activity, sensitizes cells to genotoxic damage, suggesting loss of the 5’dRP lyase activity may enhance retention of Polβ to sites of DNA damage (Sobol et al., 2000). To investigate the role of Polβ’s enzymatic activities on its recruitment and retention to DNA damage sites, EGFP-Polβ, EGFP-Polβ(K72A) and EGFP-Polβ(D256A) were expressed in U2OS cells. Surprisingly, no significant changes in recruitment kinetic profiles, time to peak intensity, or half-life of recruitment were observed (**Figures 3A-C**). As it is conceivable that endogenous Polβ was contributing to the repair of the laser-induced DNA damage, the WT and mutant fusions were modified to be gRNA resistant (using a silent mutation) and then expressed in U2OS/POLB-KO cells (**Figure 3D**). However, loss of endogenous Polβ had no effect on the recruitment profiles of either the dRP lyase or polymerase mutants (**Figure 3E**). This was unexpected, as previous studies have shown that Polβ(K72A)-expressing mouse fibroblasts are hyper-sensitive to DNA damage (Sobol et al., 2000). One explanation may be that the Polβ(K72A) mutant retains ∼1% residual 5’dRP lyase activity, which may contribute to DNA repair (Matsumoto et al., 1998). To address this, an EGFP-Polβ 5’dRP lyase triple mutant, Polβ(K35A/K68A/K72A), completely devoid of 5’dRP lyase activity (Sobol et al., 2000), was expressed in the U2OS/POLB-KO cells. The EGFP-Polβ 5’dRP lyase triple mutant did not display altered recruitment profiles as compared to EGFP-Polβ in POLB knockout cells (U2OS/POLB-KO) (**Figure 3F**). To identify if this was a U2OS cell-specific outcome, A549/POLB-KO cells were used to verify the 5’dRP lyase activity mutant results. Again, no change in recruitment was observed in the EGFP-Polβ(K72A) expressing cells compared to EGFP-Polβ (**Figures S3A-D**). In all, we find that Polβ recruitment to and retention at sites of DNA damage is not dependent on either of Polβ’s known enzymatic functions.

**Figure 3:**
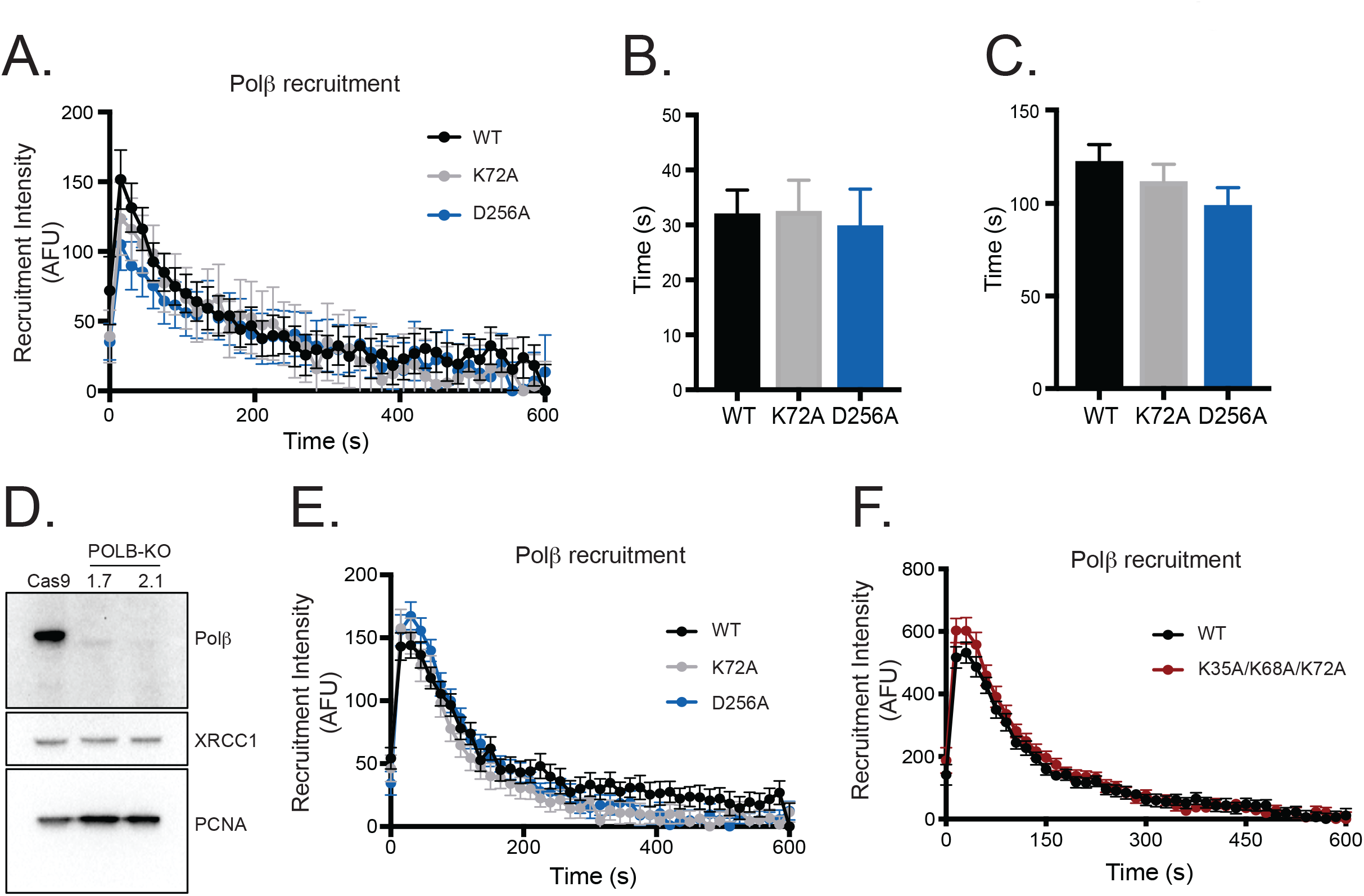
Loss of Polβ enzymatic activities does not alter its recruitment kinetics. A) Recruitment of EGFP-Polβ, dRP lyase mutant EGFP-Polβ(K72A), and polymerase mutant EGFP-Polβ(D256A) in U2OS cells following laser micro-irradiation, n ≥35. Cells retained endogenous Polβ. B) Time to peak recruitment intensity of EGFP-Polβ, EGFP-Polβ(K72A), and EGFP-Polβ(D256A) in U2OS cells following laser micro-irradiation. No significant difference was observed following a one-way ANOVA. Error bars indicate standard error of the mean. C) Half-life of recruitment of EGFP-Polβ, EGFP-Polβ(K72A), and EGFP-Polβ(D256A) following laser micro-irradiation. No significant difference was observed following a one-way ANOVA. D) Immunoblot of CRISPR/Cas9-mediated POLB KO in U2OS cells. Two cells lines were generated using two different guide RNAs. E) Recruitment of EGFP-Polβ, dRP lyase mutant EGFP-Polβ(K72A), and polymerase mutant EGFP-Polβ(D256A) in U2OS/POLB-KO(1.7) cells, n ≥35. No change in recruitment was observed when endogenous POLB was knocked out. F) Recruitment of EGFP-Polβ and dRP lyase triple mutant EGFP-Polβ(K35A/K68AK72A) in A549/POLB-KO cells following laser micro-irradiation, n ≥35. No change in recruitment, as compared to WT, was observed when using the triple mutant in the A549 cell line. See also Figure S3.

### Polβ’s recruitment is dependent on XRCC1, while Polβ enables XRCC1 complex disassociation

Recruitment to and retention of Polβ at micro-irradiation-induced DNA damage sites may be regulated through one of its binding partners. A strong candidate for this role is XRCC1, which functions as a scaffold for multiple repair proteins, including Polβ, at sites of DNA damage (Kubota et al., 1996). To investigate the functionality of this interaction in response to DNA damage, XRCC1 knockout (KO) cells were generated using CRISPR-Cas9 and verified by immunoblot (**Figure 4A, S4**). Loss of XRCC1 attenuated Polβ recruitment to sites of DNA damage in two independent KO cell lines (**Figure 4B**). This suggests a requirement for XRCC1 with regard to the recruitment and BER complex assembly for Polβ, but it does not address if physical binding is required for this effect. The V303 loop of Polβ has previously been identified as facilitating the physical interaction of Polβ with XRCC1. A separation of function mutation in this loop (i.e. L301R/V303R/V306R, referred to as Polβ(TM)) reduces the binding affinity between Polβ and XRCC1 by greater than 6-fold (Fang et al., 2019; Fang et al., 2014). EGFP-Polβ(TM) was expressed in A549 cells expressing endogenous XRCC1 and as predicted, did not visibly recruit to sites of laser micro-irradiation (**Figure 4C**), demonstrating that the physical interaction between Polβ and XRCC1 is required to facilitate Polβ recruitment.

**Figure 4:**
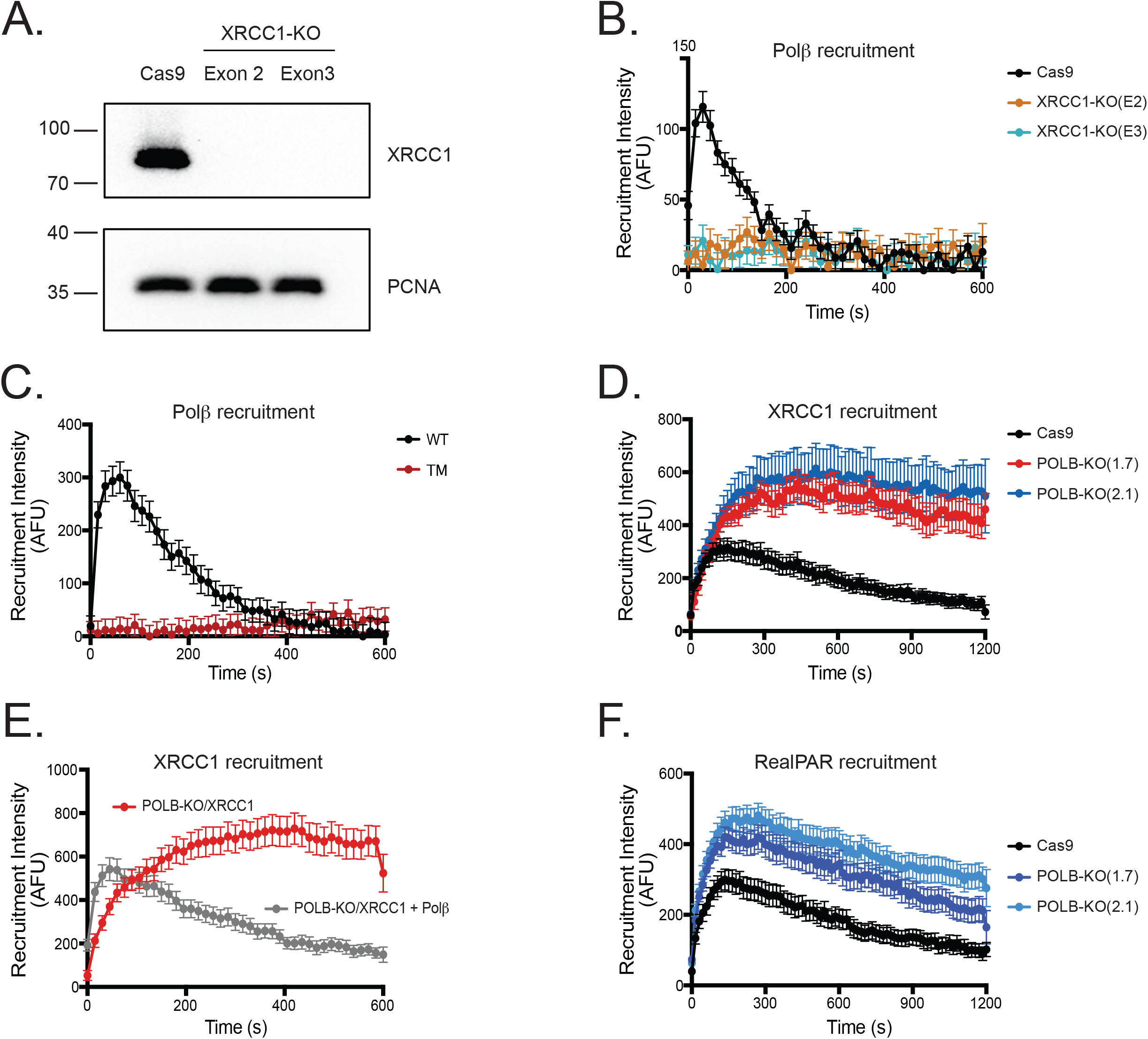
Polβ’s recruitment is dependent on XRCC1, while Polβ enables XRCC1 complex dissociation. A) Immunoblot of CRISPR/Cas9-mediated XRCC1-KO in U2OS cells. Expression of PCNA is shown as a loading control. Two cells lines were generated using two different guide RNAs. B) Recruitment of EGFP-Polβ in XRCC1-KO cells, n ≥35. Polβ failed to recruit in the absence of XRCC1. Error bars indicate standard error of the mean. C) Recruitment of EGFP-Polβ and XRCC1-binding deficient triple mutant EGFP-Polβ (L301R/V303R/V306R) in A549 cells following laser micro-irradiation, n ≥35. Polβ triple mutant failed to recruit, even with endogenous XRCC1 expressed. D) Recruitment of XRCC1-EGFP in U2OS/POLB-KO cells following laser micro-irradiation, n ≥35. XRCC1 demonstrated the ability to recruit to micro-irradiation-induced DNA damage in the absence of Polβ, but XRCC1 demonstrated slower disassembly from the repair complex in the absence of Polβ. E) Recruitment of XRCC1-EGFP in U2OS/POLB-KO cells with POLB restored, n ≥35. XRCC1 demonstrated slower disassembly from the repair complex in the absence of Polβ. Re-expression of Polβ was able to restore the disassembly of XRCC1 from sites of DNA damage. F) Recruitment of RealPAR in U2OS/POLB-KO cells, n ≥35. U2OS/POLB-KO cells following laser micro-irradiation demonstrated increased PAR formation in comparison to U2OS cells but no change in PAR degradation in the absence of POLB. See also Figure S4.

Because the recruitment of XRCC1 to the site of damage was critical for the recruitment of Polβ, we then investigated if alterations in Polβ could modulate XRCC1 recruitment. XRCC1-EGFP recruitment exhibited both enhanced peak recruitment intensity and prolonged recruitment in POLB-KO cells when compared to Polβ-expressing cells (**Figure 4D**). This novel finding suggests that Polβ is required to facilitate XRCC1 dissociation from assembled DNA repair complexes. To validate if this was the case, Polβ and XRCC1-EGFP were co-expressed in POLB-KO cells. By re-expressing Polβ, we were able to recapitulate XRCC1’s disassociation from micro-irradiation-induced foci, thereby demonstrating a requirement for Polβ to promote rapid dissociation of XRCC1 from the site of DNA damage (**Figure 4E**). These results suggest a negative feedback loop in which Polβ requires XRCC1 for recruitment to sites of DNA damage, but then itself acts as a regulator of XRCC1 dissociation from those sites.

The dependence of XRCC1 on Polβ for dissociation may be mediated by enhanced PARylation at the site of DNA damage in the absence of Polβ, as Polβ loss would likely lead to prolonged PAR formation at the site of damage (Jelezcova et al., 2010; Tang et al., 2010). To determine if this was caused by altered PAR formation, we utilized the RealPAR probe to interrogate PAR dynamics in POLB-KO cells. PARylation was enhanced in POLB-KO cells compared to normal cells, but the rate of PAR degradation, as inferred from the change in RealPAR binding, was similar in POLB-KO cells compared to Polβ expressing cells (**Figure 4F**), suggesting that the requirement of Polβ for XRCC1’s rapid complex disassociation is not dependent on PAR formation.

### Polβ and XRCC1 complex dynamics depend on PAR formation and degradation

PARylation is a critical component of BER/SSBR (Dantzer et al., 2000; Schreiber et al., 2002), so we investigated how PARylation alters Polβ/XRCC1 repair complex assembly. To identify how PARP1 regulates Polβ complex assembly, PARP1 knockout cells were generated and validated by immunoblot (**Figure 5A**). Polβ recruitment was attenuated but not eliminated in PARP1 knockout cells, which is consistent with previous findings that other PARPs (such as PARP2) promote PARylation and DNA repair complex assembly during BER (**Figure 5B**) (Dantzer et al., 2000; Schreiber et al., 2002). To address the role of PARylation in Polβ/XRCC1/PAR repair complex assembly and disassembly more broadly, the PARP inhibitor ABT-888 (velipirab) and the PARG inhibitor PDD00017273 were used. As shown above, PARylation, as determined by the recruitment of the RealPAR probe, was abolished with ABT-888 treatment while PARG inhibition enhanced and prolonged PAR formation at sites of laser micro-irradiation (**Figure 1G**). Polβ and XRCC1 behaved similarly, showing attenuation of recruitment upon PARP inhibition and prolonged recruitment and retention following PARG inhibition (**Figures 5C-D**). PARylation synthesis and degradation therefore temporally regulate Polβ and XRCC1 repair complex assembly and disassembly at sites of DNA damage. In addition to PARG’s degradation of PAR from sites of DNA damage, TARG (OARD1) serves to degrade PAR by removing the O-acyl-ADP-ribose moiety that is the first ADP-ribose added to a PAR chain (Sharifi et al., 2013). To determine if TARG impacts Polβ complex formation, TARG was knocked out using CRISPR/Cas9 (**Figure S5A**), but no change in Polβ recruitment kinetics was observed (**Figure S5B**).

**Figure 5:**
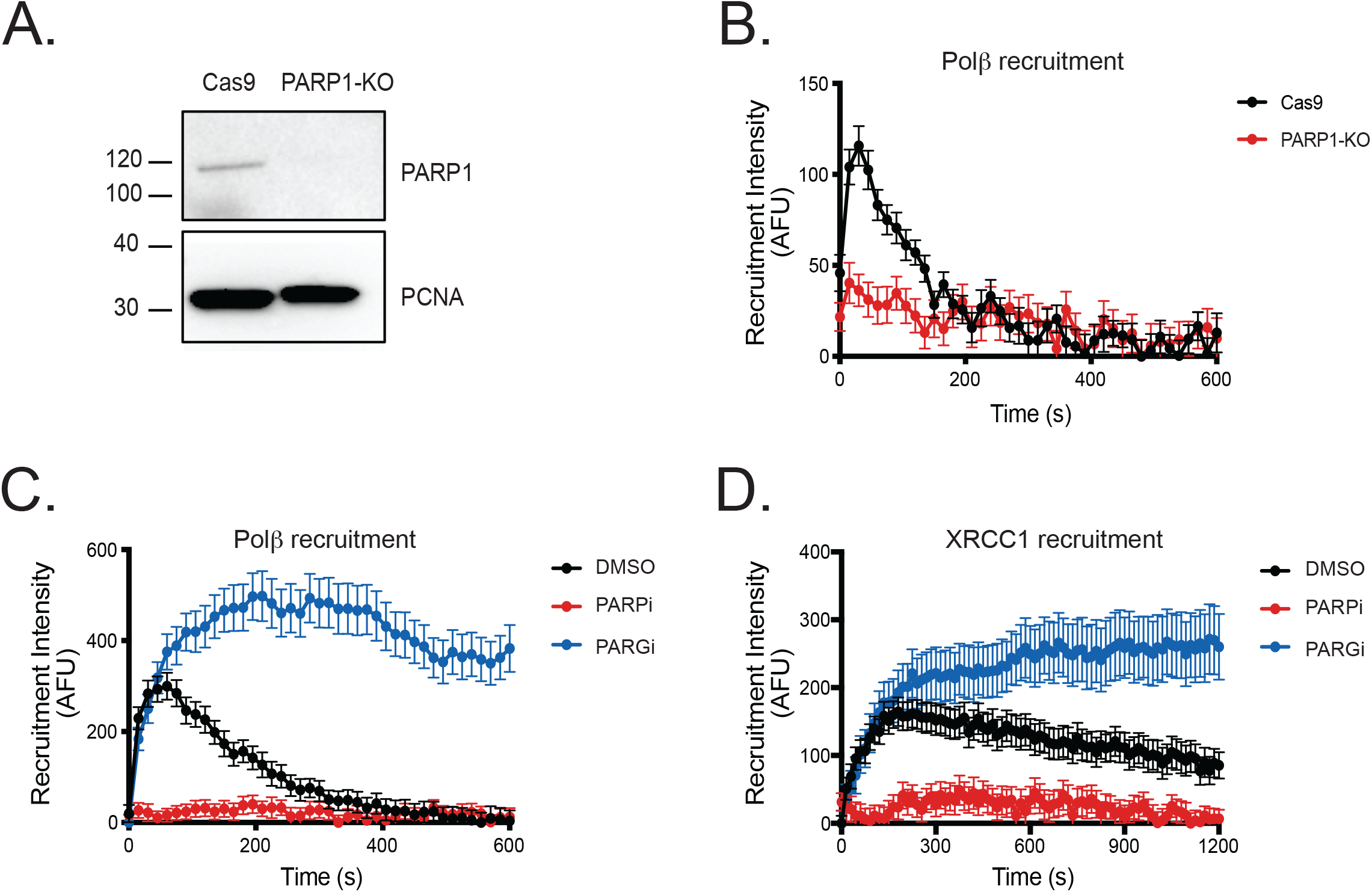
Polβ and XRCC1 complex dynamics are dependent on PAR formation and degradation. A) Immunoblot of CRISPR/Cas9-mediated PARP1-KO in U2OS cells. B) Recruitment of EGFP-Polβ in U2OS/PARP1-KO cells following laser micro-irradiation, n ≥35. Polβ recruitment was attenuated, in the absence of PARP1. Error bars indicate standard error of the mean. C) Recruitment of EGFP-Polβ in A549 cells following laser micro-irradiation with PARP or PARG inhibition, n ≥35. PARP inhibition (10μM ABT888) attenuated Polβ recruitment, and PARG inhibition (10μM PDD00017273) enhanced Polβ’s peak recruitment intensity and prolonged focal recruitment at sites of micro-irradiation. D) Recruitment of XRCC1-EGFP in A549 cells following laser micro-irradiation with following PARP or PARG inhibition, n ≥35. PARP inhibition attenuated XRCC1 recruitment, and PARG inhibition enhanced XRCC1’s peak recruitment intensity and prolonged focal recruitment at sites of micro-irradiation. See also Figure S5.

### Polβ and XRCC1 complex assembly is regulated by NAD^+^ availability

NAD^+^ is a metabolic product and regulator that is an essential substrate for PARPs to produce PAR chains (Fouquerel and Sobol, 2014; Rouleau et al., 2010). Because PAR is necessary for Polβ and XRCC1 recruitment to DNA damage, modulation of NAD^+^ should lead to changes in Polβ and XRCC1 recruitment. To identify how NAD^+^ availability affects Polβ/XRCC1 repair complex formation, the NAMPT inhibitor FK866 was used to decrease cellular NAD^+^ levels, and dihydronicotinamide riboside (NRH) was used to enhance cellular NAD^+^ levels. Due to factors in serum that alter NAD^+^ catabolite stability (Wilk et al., 2020), we used heat-inactivated fetal bovine serum (HI-FBS) for these studies. We noted that the recruitment profiles of Polβ, XRCC1, or RealPAR with HI-FBS, as compared to normal fetal bovine serum, were unchanged (**Figures S6A-C**).

FK866 treatment diminished NAD^+^ levels in U2OS cells to 23% of control (**Figure 6A**). Since the conversion of NRH to NAD^+^ is dependent on the metabolic profile of the cell, with maximal NAD^+^ enhancement observed at different times in different cells (Giroud-Gerbetant et al., 2019; Yang et al., 2019), a time-course was performed in U2OS cells to identify when NAD^+^ levels were maximally enhanced following NRH administration (**Figure 6B**). A peak in NAD^+^ concentration was observed at 4 hours post-NRH addition (∼850% increase), with NAD^+^ diminishing but remaining above controls for up to 8 hours post-treatment. NRH also enhanced PARP1 activation capacity and DNA damage-induced (H_2_O_2_) PARylation in U2OS cells, an effect attenuated by FK866 or ABT888 treatment (**Figure 6C**). In laser micro-irradiation experiments, NRH enhanced peak recruitment intensities of Polβ (45%), XRCC1 (94%), and RealPAR (88%) (**Figures 6D-F**). Conversely, FK866 reduced peak recruitment intensities of Polβ (37% of control), XRCC1 (35%), and RealPAR (24%) (**Figure 6G-I**). No change in the time to peak recruitment intensity or half-life of recruitment was observed, suggesting that NAD^+^ availability does not impact repair complex assembly or disassembly other than to increase the magnitude of the recruitment.

**Figure 6:**
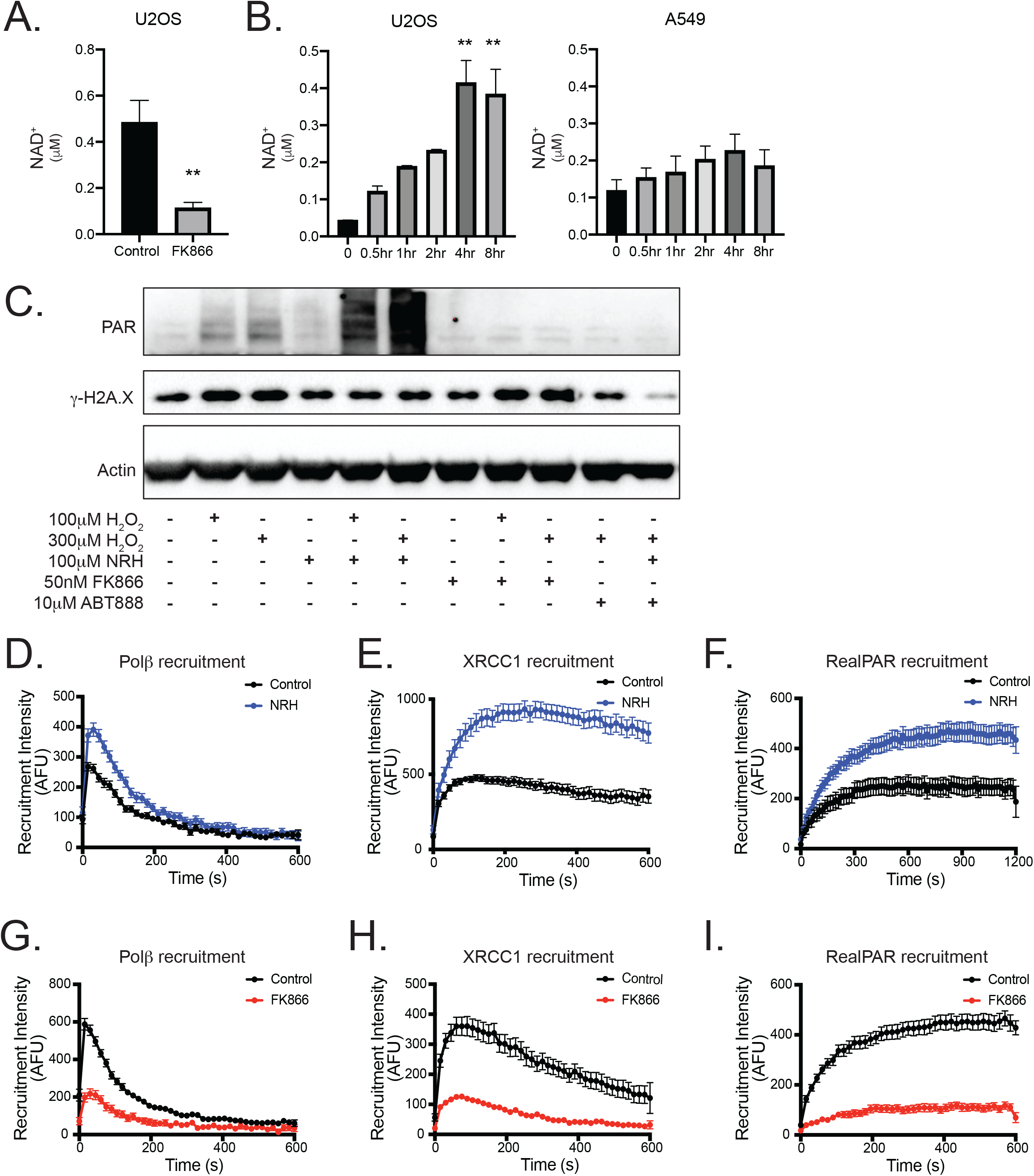
Polβ and XRCC1 complex dynamics are regulated by NAD^+^ bioavailability. A) NAD^+^ concentrations in U2OS cells following treatment with the NAMPT-inhibitor FK866 (10μM, 24 hours). FK866 reduced NAD^+^ concentrations to 23% of control (p<0.01; Student’s t-test). Graphs depict mean and standard error of the mean. B) Time course of NAD^+^ concentrations in U2OS cells following NRH treatment (100μM). NRH increased NAD^+^ concentrations by 850% in U2OS cells following 4 hours of treatment (p<0.01; one-way ANOVA and a Tukey *post-hoc* test). NRH did not enhance NAD^+^ in A549 cells. Graphs depict mean and standard error of the mean. C) Immunoblots revealing PAR formation in U2OS cells following a combination of H_2_O_2_, NRH, FK866, or ABT-888 treatment. PAR formation was stimulated using H_2_O_2_ (100μM or 300μM). NRH enhanced PARylation following H_2_O_2_ treatment, while ABT-888 was able to block the effect. Similarly, FK866 pre-treatment attenuated PAR formation upon H_2_O_2_ treatment. D) Recruitment of EGFP-Polβ to sites of laser micro-irradiation (355nm) in U2OS cells following NRH treatment, n ≥35. Polβ peak recruitment intensity was enhanced (45%) following 4 hours NRH pre-treatment (100μM). Error bars indicate standard error of the mean. E) Recruitment of XRCC1-EGFP to sites of laser micro-irradiation (355nm) in U2OS cells following NRH treatment, n ≥35. XRCC1 peak recruitment intensity was enhanced (94%) following 4 hours NRH pre-treatment. F) Recruitment of RealPAR to sites of laser micro-irradiation (355nm) in U2OS cells following NRH treatment, n ≥35. RealPAR peak recruitment intensity was enhanced (88%) following 4 hours NRH pre-treatment. G) Recruitment of EGFP-Polβ to sites of laser micro-irradiation (355nm) in U2OS cells following FK866 treatment (10μM), n ≥35. Error bars indicate standard error of the mean. H) Recruitment of XRCC1-EGFP to sites of laser micro-irradiation (355nm) in U2OS cells following FK866 treatment, n ≥35. I) Recruitment of RealPAR to sites of laser micro-irradiation (355nm) in U2OS cells following FK866 treatment, n ≥35. See also Figure S6.

To determine the broad applicability of NRH for enhancing intracellular NAD^+^, the same assays were also performed in A549 cells. Interestingly, NRH did not significantly increase NAD^+^ in A549 cells (**Figure 6B**), possibly due to the low expression of adenosine kinase in this cell line (Protein-Atlas, 2020; Uhlen et al., 2017; Yang et al., 2020). It is possible that NAD^+^ formed via NRH metabolism was being converted into NADH in A549 cells, but we found no changes in NADH following NRH supplementation, in either A549 or U2OS cells (**Figure S6D**). Utilization of U2OS and A549 cell lines provided a unique opportunity to identify effects (if any) of NRH on DNA repair complex assembly independent of its NAD^+^-enhancing capability. Following NRH treatment (100μM, 4hrs), A549 did not demonstrate enhanced Polβ, XRCC1, or RealPAR recruitment kinetics (**Figures S6E-G**). In all, we find that NRH’s ability to enhance NAD^+^ is dependent on the ability of a cell to metabolize NRH into NAD^+^, and NRH’s enhancement of NAD^+^ directly impacts damage-induced PAR synthesis and the recruitment of Polβ, XRCC1, and RealPAR to sites of DNA damage.

### Loss of SIRT6 impairs Polβ and XRCC1 complex assembly without altering PAR formation

SIRT6 was previously documented to play a critical and essential role in BER mediated repair but the exact mechanism has never been resolved. Loss of SIRT6 enhanced genomic instability caused by alkylation and oxidation damage, and supplementation of SIRT6-deficient cells with Polβ’s dRP lyase domain was able to protect against SIRT6-KO genotoxic sensitivity (Mostoslavsky et al., 2006). It was suggested that SIRT6 may modulate repair by regulating PARP1 (Mao et al., 2011) such that SIRT6 is required for PARP1 activation in DSB repair (Tian et al., 2019). We therefore utilized RealPAR together with the Polβ and XRCC1 fusion proteins to evaluate the role of SIRT6 on Polβ and XRCC1 recruitment and PAR formation in response to DNA damage in WT and SIRT6-KO cells (**Figure 7A**). Both Polβ and XRCC1 demonstrated significantly diminished recruitment to DNA damage in SIRT6 knockout cells (**Figures 7B-C**). However, loss of SIRT6 did not lead to a change in RealPAR focal recruitment, demonstrating that PAR formation was not altered in SIRT6-KO cells (**Figure 7D**). Together, these results support a model where SIRT6 enhances XRCC1’s binding to PAR at sites of DNA damage, and loss of SIRT6 diminishes XRCC1 (and by extension Polβ) recruitment to sites of DNA damage (**Figure 7E**).

**Figure 7:**
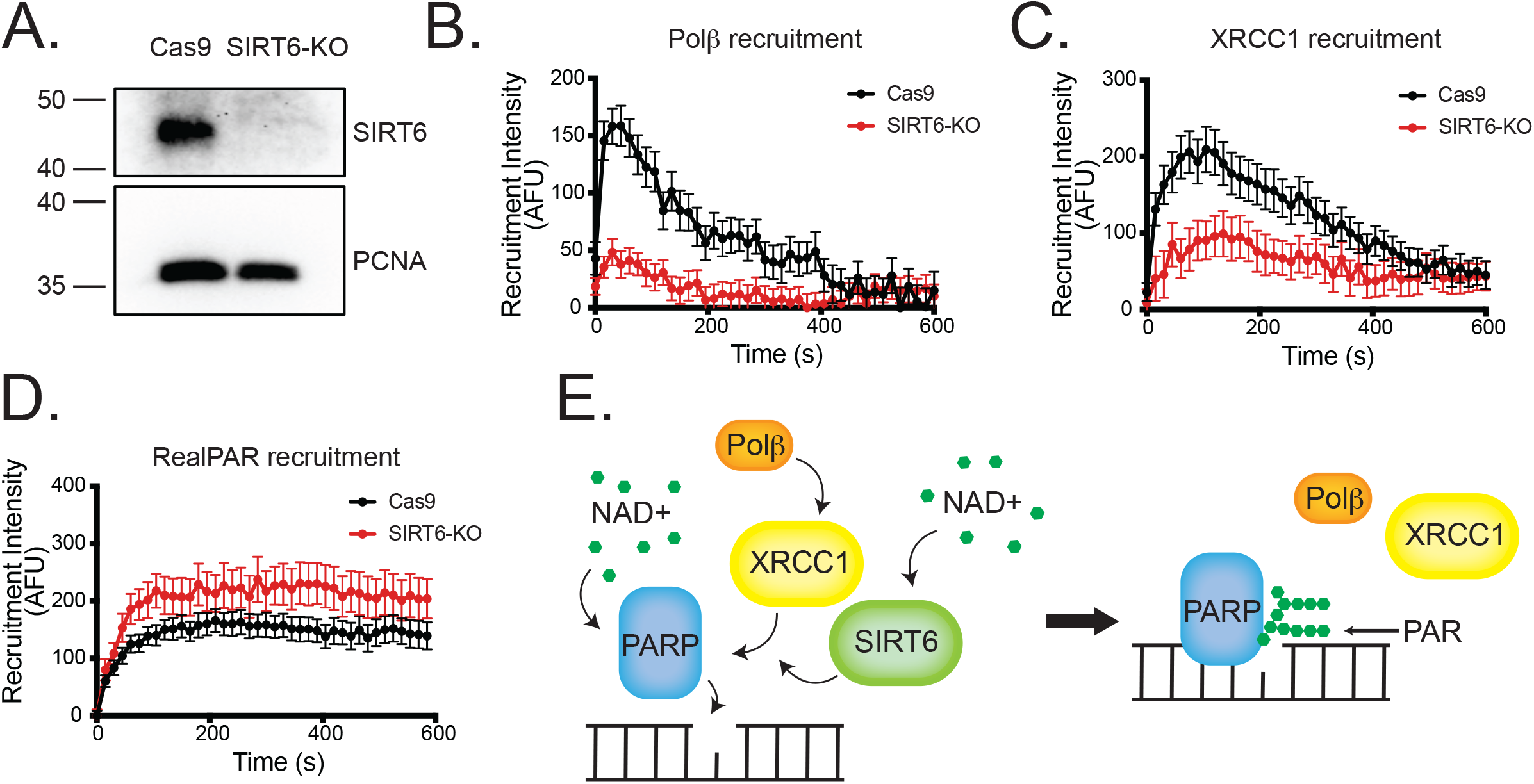
Loss of SIRT6 impairs Polβ and XRCC1 complex assembly without altering PAR formation. A) Immunoblot of CRISPR/Cas9-mediated SIRT6-KO in A549 cells. B) Recruitment of EGFP-Polβ to sites of laser micro-irradiation (355nm) in SIRT6-KO cells, n ≥35. Polβ recruitment was attenuated in the absence of SIRT6. Error bars indicate standard error of the mean. C) Recruitment of XRCC1-EGFP to sites of laser micro-irradiation (355nm) in SIRT6-KO cells, n ≥35. XRCC1 recruitment was attenuated in the absence of SIRT6. D) Recruitment of RealPAR to sites of laser micro-irradiation (355nm) in SIRT6-KO cells, n ≥35. RealPAR recruitment was unchanged in the absence of SIRT6. E) Model of SIRT6 activity on PAR-dependent recruitment of XRCC1 to sites of DNA damage. SIRT6 regulates the ability of XRCC1 to bind to PAR following micro-irradiation. PAR formation is unchanged in SIRT6-KO cells. The reduction in SIRT6 reduces the recruitment of XRCC1 binding proteins, including Polβ.

## DISCUSSION

The characterization of DNA repair complex assembly and disassembly remains an intensive area of investigation for multiple biological fields, including cancer and aging. Identification of key substrates, enzymatic proteins, and accessory factors are key in refining our models of DNA repair pathway control and cross-talk. Here, we investigated the formation of BER/SSBR complex assembly and disassembly in live cells using laser micro-irradiation. While the sequence of enzymatic steps required to repair BER/SSBR lesions has been biochemically characterized using *in vitro* assays, laser micro-irradiation of fluorescently tagged repair proteins enables interrogation of the repair process within an intact cellular context, allowing analysis of the effects of loss or modulation of key DNA repair components. We used the novel MIDAS system to acquire time-lapse images of laser-induced fusion protein recruitment, detection and quantitation of repair foci in resulting images, and analysis of recruitment kinetics for key parameters.

We first identified the recruitment kinetics of Polβ and XRCC1, confirming that Polβ’s recruitment was attenuated in XRCC1 knockout cells and was dependent on its V303 loop as previously described (**Figure 4C**) (Fang et al., 2014). Interestingly, we observed Polβ dissociating from the repair complex before XRCC1 (**Figures 1B-C**). This was unexpected because Polβ and XRCC1 are thought to form a heterodimer during repair and it was anticipated that recruitment profiles of Polβ and XRCC1 should be similar (Fang et al., 2014). We considered that over-expression of the enzymatic component of the repair complex (EGFP-Polβ) might enable faster repair of the DNA damage, as compared to cells with lower levels of the EGFP-Polβ fusion protein. This might then explain why EGFP-Polβ over-expressing cells were displaying faster recruitment kinetics, but experiments using endogenously tagged Polβ demonstrated that this was not the case (**Figure 2**). It is currently unknown if Polβ and XRCC1 are recruited as a heterodimer to sites of DNA damage or if they combine at the site, but we observed no difference in XRCC1’s ability to assemble at sites of DNA damage when Polβ was knocked out (**Figure 4D**). In either case, the results suggest that Polβ is being removed from the site prior to XRCC1, which implicates an additional control mechanism for Polβ repair complex disassembly.

Because our model predicted that XRCC1 recruitment was dependent on PAR but not on Polβ, we investigated if loss of Polβ affected XRCC1 recruitment. Our results showed that loss of Polβ enhanced XRCC1 recruitment but also increased XRCC1 retention to sites of laser-induced damage (**Figure 4D**). Conversely, re-expression of Polβ in POLB knockout cells restored XRCC1’s disassembly kinetics (**Figure 4E**). This was a novel finding but could be explained by incomplete repair of the damage site due to a lack of Polβ’s enzymatic activities. However, neither EGFP-tagged mutants for Polβ’s dRP lyase (K72A or K35A/K68A/K72A) nor polymerase (D256A) activities demonstrated altered recruitment kinetics compared to functional Polβ (**Figures 3A**), even when endogenous Polβ was removed (**Figures 3E-F**). These results have two implications. First, the results identify a novel non-enzymatic function of Polβ in facilitating XRCC1 repair complex disassembly. Second, it shows that neither the dRP lyase or polymerase activity of Polβ is required for assembly or disassembly of Polβ at repair foci. Together, these results support a model where Polβ’s recruitment to and dissociation from repair complexes is not dependent on repair of the DNA lesion itself (or repair is facilitated by compensatory enzymes), and that Polβ further facilitates removal of XRCC1 from the DNA damage sites. This model is further supported by the lack of change observed in assembly/disassembly kinetics between EGFP-Polβ over-expression and endogenously expressed EGFP-Polβ (**Figure 2E**).

Due to the dependence of Polβ and XRCC1 on PAR formation for repair, we investigated the role of PAR on recruitment dynamics. Previous micro-irradiation research has utilized PAR antibodies to characterize PAR formation in cells. The results can be impacted by the fixation method used, the antibody used and is hampered by the loss of temporal resolution (a key concern for short time frames). To address this limitation, we developed the novel RealPAR probe to provide real-time, live-cell imaging of PAR formation at sites of laser micro-irradiation (**Figure 1D-E**). RealPAR’s focal recruitment is dependent on its ability to bind PAR and as we show, does not require BrdU or Hoechst sensitization for recruitment visualization using either 355nm and 405nm laser wavelengths. Further, we show that RealPAR is responsive at micro-irradiation settings that elicit Polβ and XRCC1 recruitment, two factors whose recruitment is dependent on PAR formation. We confirmed that PARP inhibition prevented PAR formation at sites of laser-induced DNA damage and was required for Polβ and XRCC1 recruitment, while loss of PAR degradation enhanced retention of Polβ and XRCC1 at sites of damage (**Figures 5C-D**). These experiments confirmed that the presence of PAR at sites of DNA damage is a critical step for Polβ and XRCC1 recruitment.

NAD^+^ supports DNA repair by serving as a substrate for both PARPs and sirtuins. Depleted NAD^+^ intracellular pools have been linked to decreased genomic integrity, and attempts to boost NAD^+^ as a mechanism for promoting genomic integrity remains active pursuits in the aging field (Fang et al., 2017). Here, we utilized NRH to promote intracellular NAD^+^ pools. NRH has been shown to enhance NAD^+^ in multiple cell types, with greater increases in NAD^+^ as compared to other NAD^+^ supplements (Giroud-Gerbetant et al., 2019; Yang et al., 2019). We demonstrated that NRH supplementation enhances intracellular NAD^+^ in U2OS cells, leading to increased assembly of Polβ/XRCC1 repair complexes (**Figure 6**). Interestingly, NRH was unable to enhance NAD^+^ in A549 cells. While the complete mechanism of NRH metabolism is unknown, adenosine kinase (ADK or AK) has been identified as one of the enzymes for NRH conversion to NAD^+^, and differential cell expression of ADK can alter NRH metabolism (Yang et al., 2020).

Finally, we utilized our Polβ/XRCC1/RealPAR system to characterize how deficiency of SIRT6 leads to compromised BER. Loss of SIRT6 enhances cellular sensitivity to genotoxins, while enhancing Polβ’s dRP lyase activity attenuates this sensitivity, demonstrating SIRT6’s involvement in Polβ-mediated repair (Mostoslavsky et al., 2006). Our Polβ/XRCC1/RealPAR system provides a unique opportunity to investigate this issue. We found that SIRT6 knockout does not alter PAR formation following micro-irradiation while XRCC1 recruitment is attenuated (**Figure 6D**). These results directly implicate SIRT6 in enhancing XRCC1’s localization to sites of damage, likely diminishing XRCC1’s ability to bind to PAR or be retained at PAR sites. Due to Polβ’s dependence on XRCC1 for repair complex assembly, it was predicted that Polβ’s recruitment to sites of DNA damage would also diminish, which we demonstrated (**Figure 6B**). This result would explain the observations from previous studies where enhancing Polβ dRP lyase activity reduces genotoxic sensitivity to SIRT6 loss (Mostoslavsky et al., 2006). Due to the importance of XRCC1 to serve as a scaffold for multiple DNA repair enzymes (e.g. Polβ, PNKP, APLF, APTX, LigIII) at sites of DNA damage, these results suggest that SIRT6 negatively impacts multiple DNA repair enzymes in BER and SSBR.

## ACKNOWLEDGMENTS

RWS is an Abraham A. Mitchell Distinguished Investigator. Research in the Sobol lab on DNA repair, the analysis of DNA damage and the impact of genotoxic exposure is funded by grants from the National Institutes of Health (NIH) [CA148629, ES014811, ES029518, ES028949 and CA238061], from the National Science Foundation (NSF) [NSF-1841811] and a grant from the DOD [GRANT11998991, DURIP-Navy]. Support is also provided from the Abraham A. Mitchell Distinguished Investigator Fund and from the Mitchell Cancer Institute Molecular & Metabolic Oncology Program Develop fund (to RWS). Support for the development of RealPAR was also provided by the Mitchell Cancer Institute Junior Faculty Award (to CAK). We thank Dr. Natalie Gassman (USA Mitchell Cancer Institute) for her invaluable conceptual input on the development of the MIDAS software platform.

## AUTHOR CONTRIBUTIONS

C.A.K., K.M.S., and R.W.S. designed the experiments. C.A.K., K.M.S, J.A., J.C., Q.F., J.L., conducted the experiments, J.F.A. developed software (MIDAS) and M.V.M., M.M. synthesized and provided critical reagents. C.A.K. and R.W.S wrote the paper. All authors read, commented on, and approved the final manuscript version.

## CONFLICT OF INTEREST

RWS is a scientific consultant for Canal House Biosciences, LLC. The authors state that there is no conflict of interest.

## Methods

### Key Resources Table

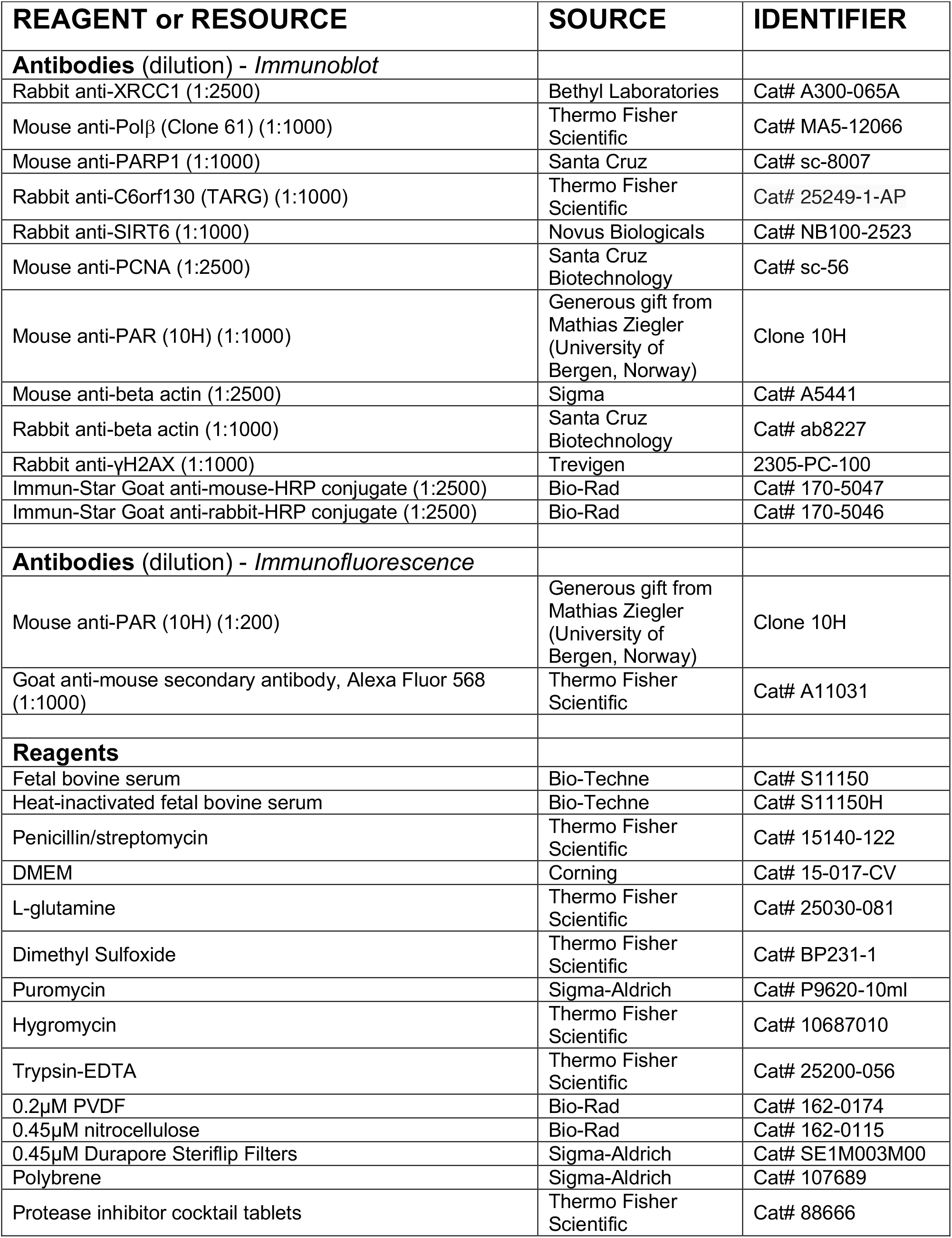
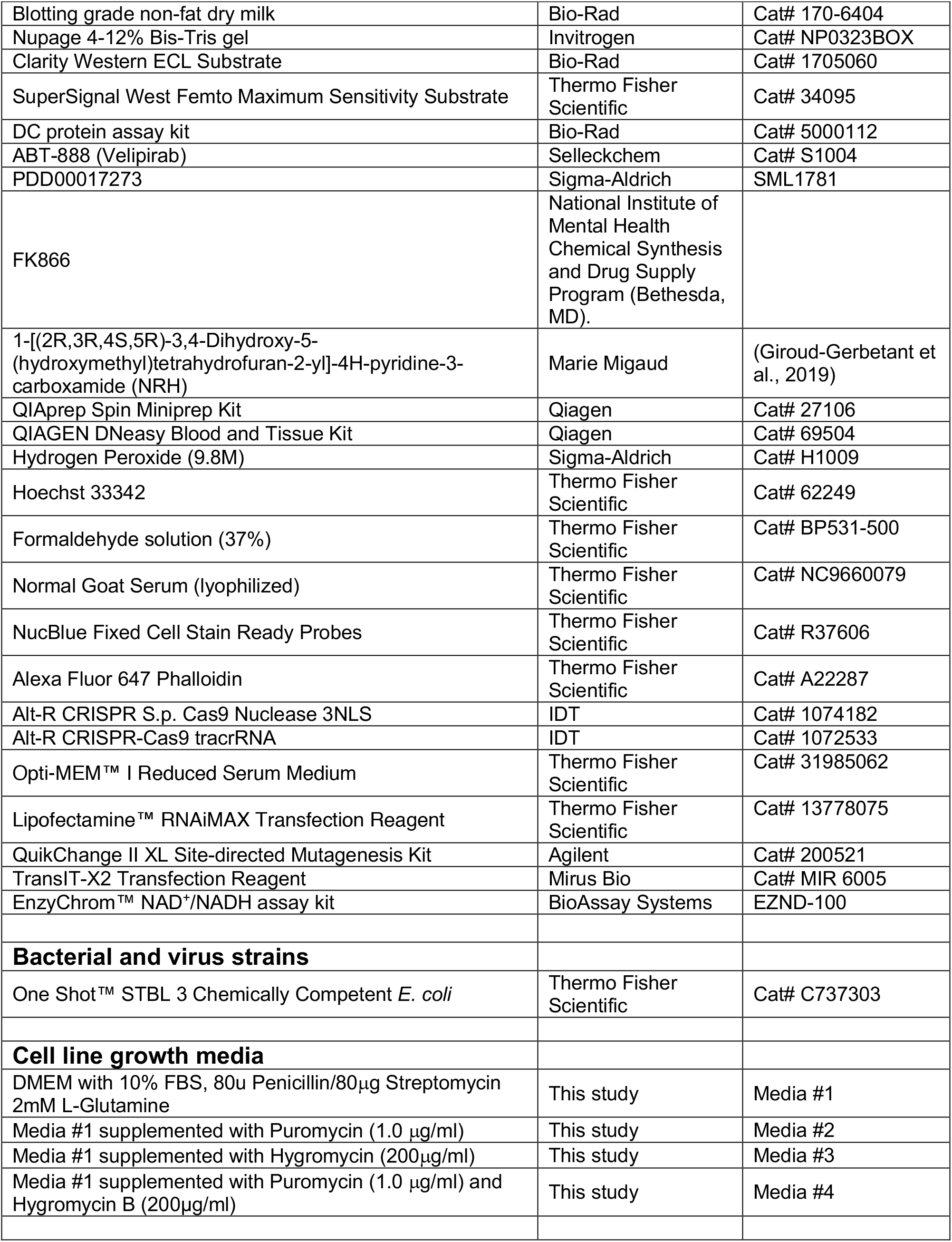
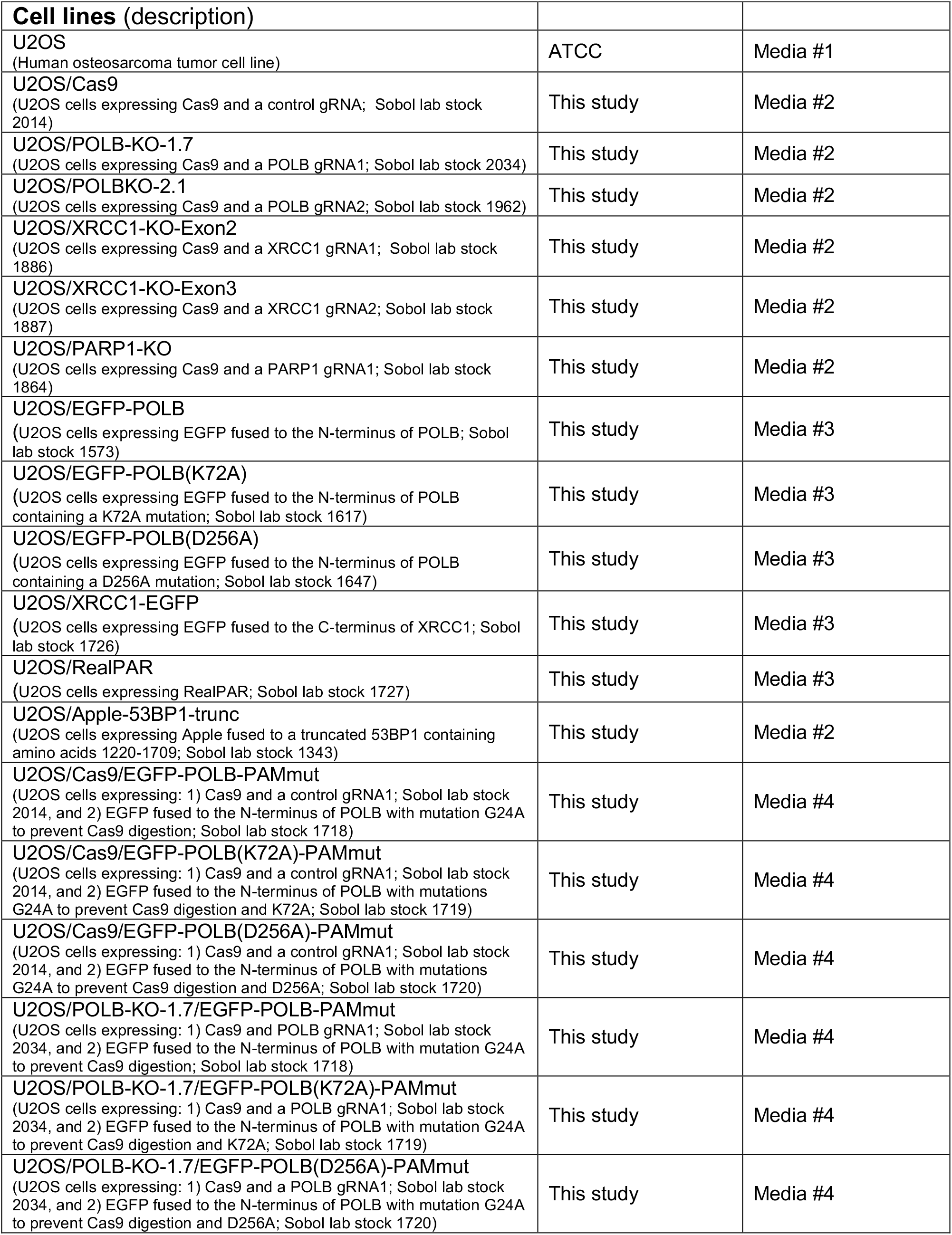
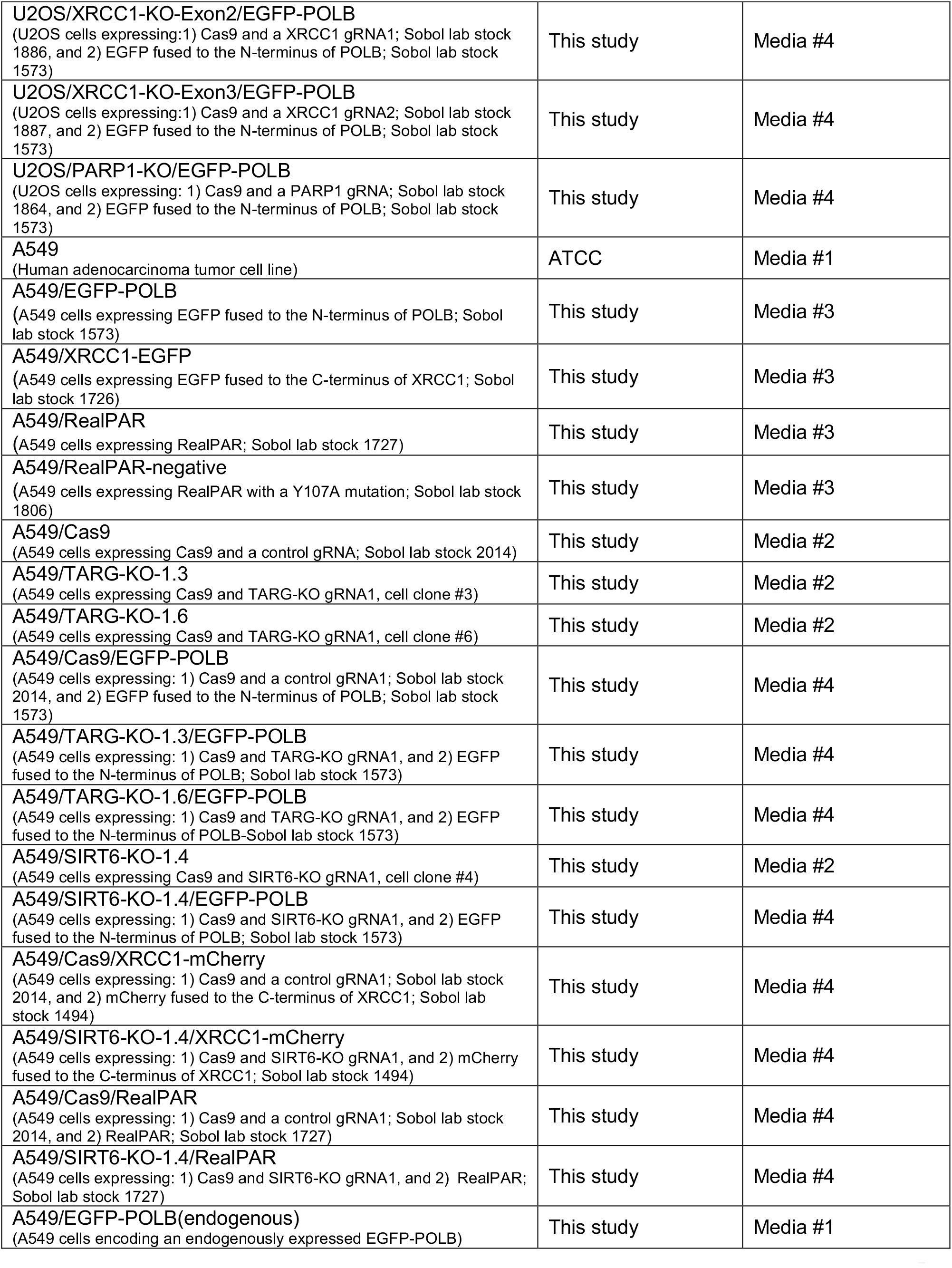
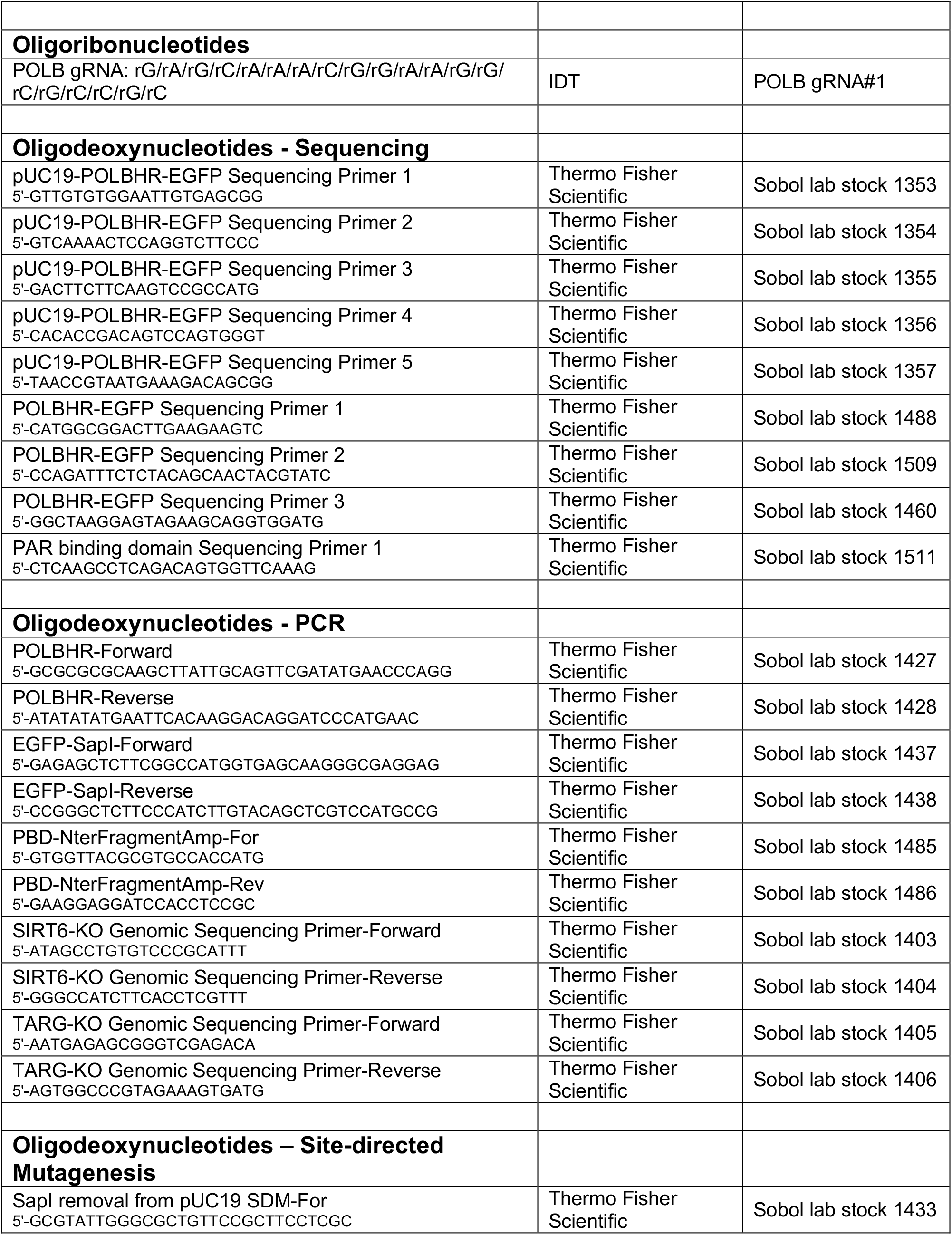
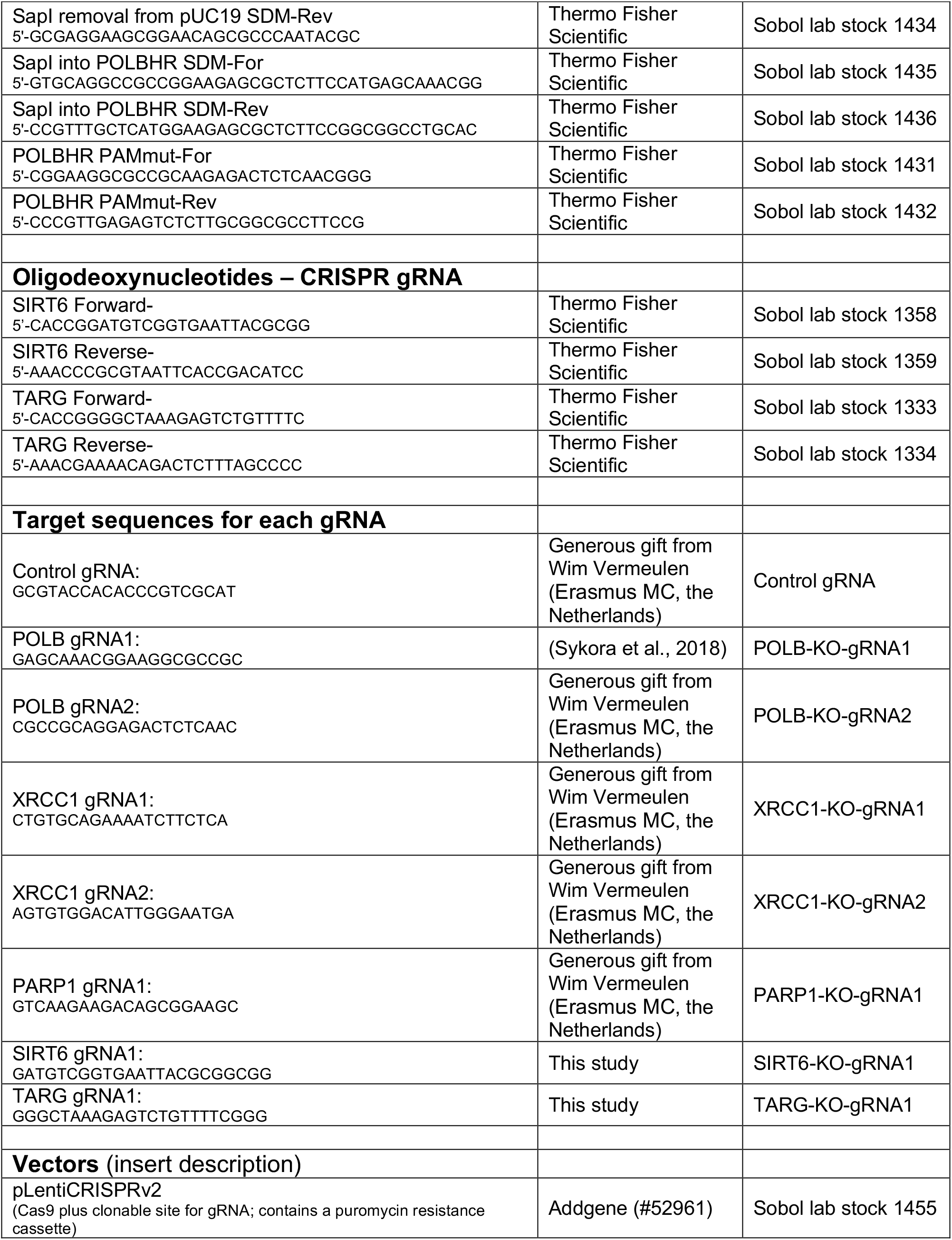
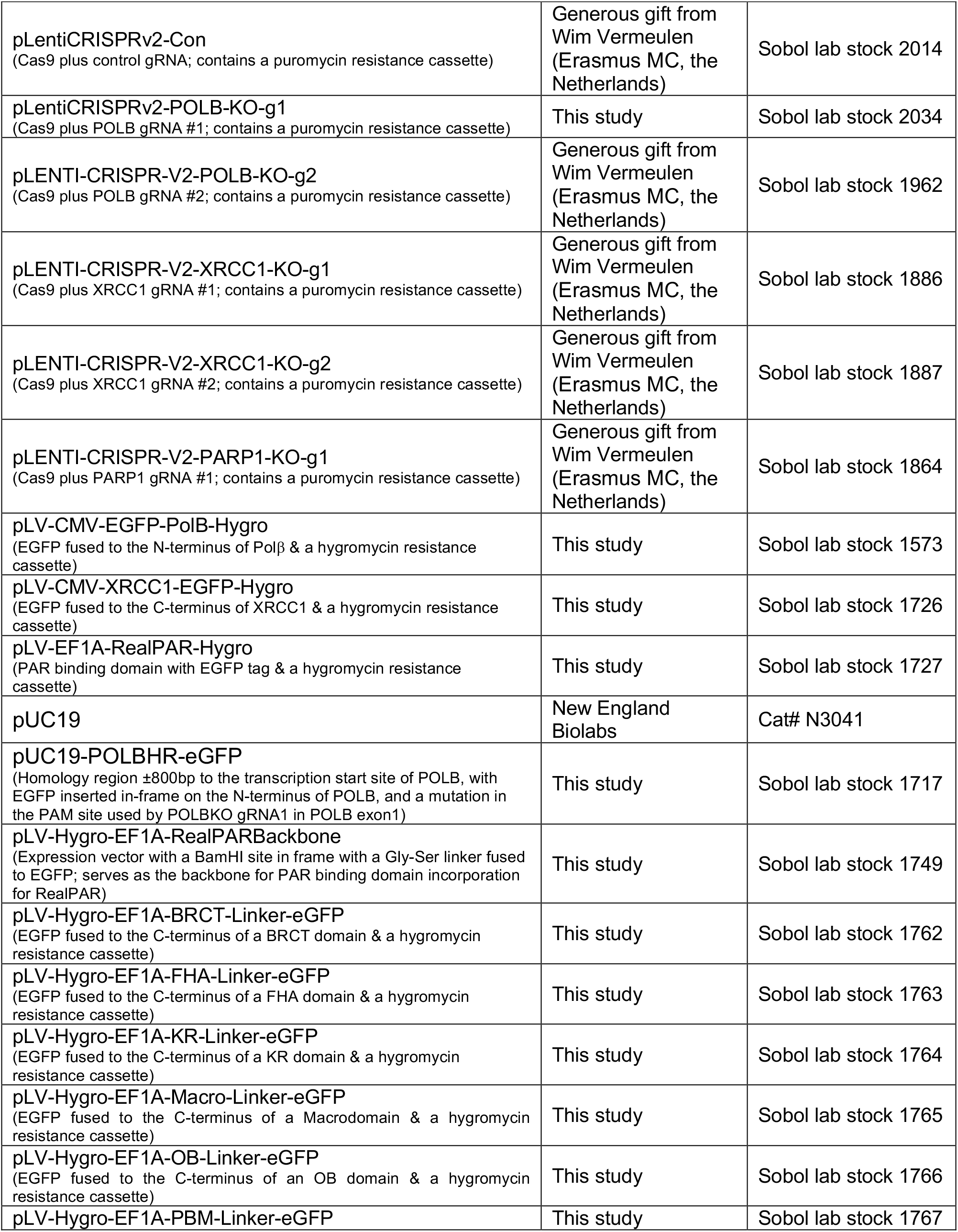
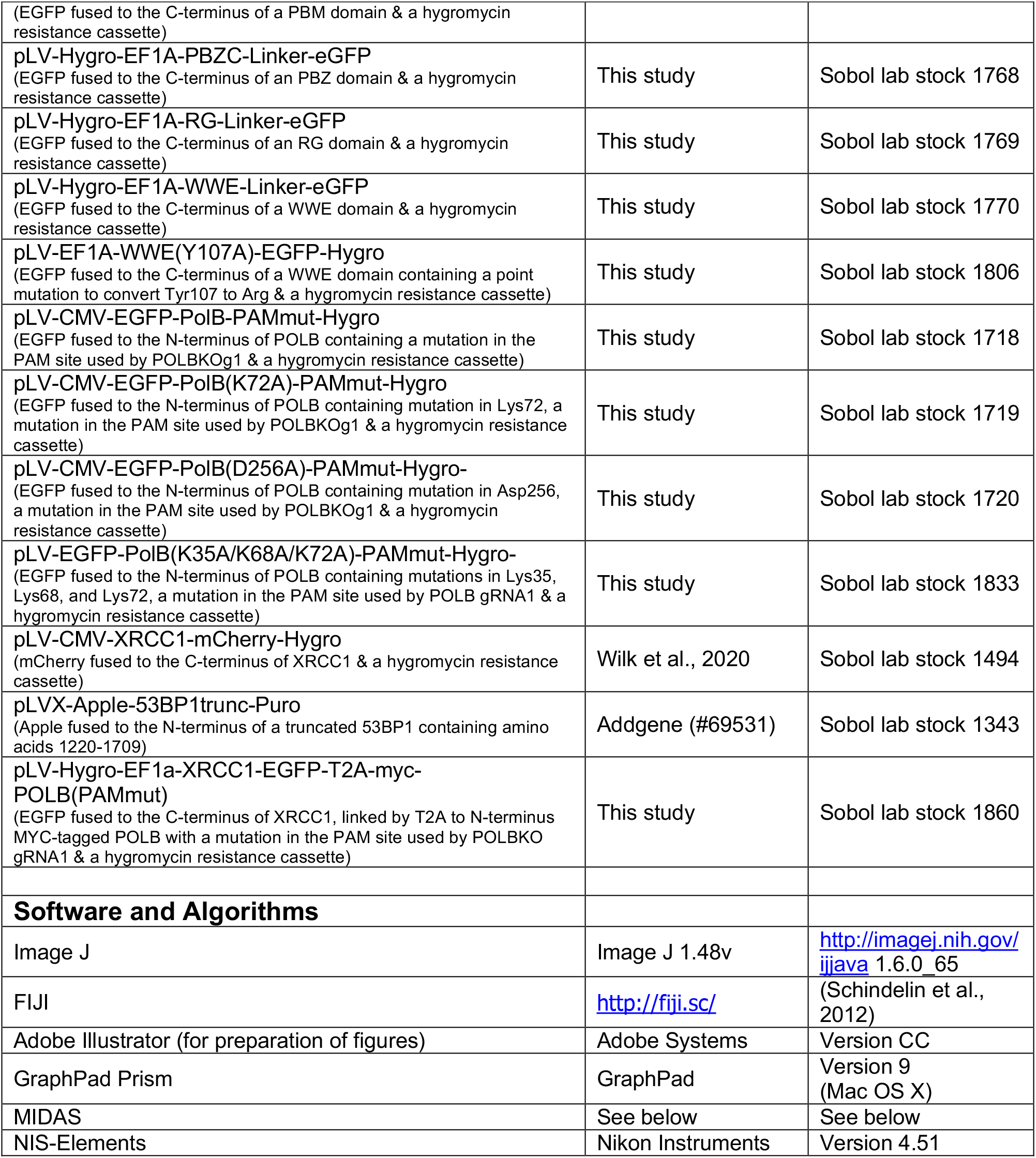

### Contact for reagent and resource sharing

Further information and requests for resources and reagents, including custom analysis scripts, should be directed to and will be fulfilled by Robert W. Sobol.

### Experimental Model and Subject Details

The human tumor cell lines A549 and U2OS were obtained from ATCC and are routinely validated by Genetica Cell Line Testing. Human tumor cell lines were modified by lentiviral-mediated expression of the indicated proteins as indicated below and as listed in the **Key Resources Table** above. In some cases, modified cells were iteratively modified by a second transduction, such as expressing POLB-KO-gRNA in cells followed by expression of EGFP-Polβ-PAMMut using lentiviral vectors with different selection makers (Puromycin, Hygromycin). Lentiviral constructs containing EGFP or mCherry fused to POLB, XRCC1 and PAR binding domains were generated for visualization of protein recruitment to sites of laser-induced (micro-irradiation) DNA damage. Genomic modification of A549 cells to introduce EGFP into the endogenous POLB gene was used for visualization of Polβ protein recruitment with laser-induced micro-irradiation experiments when evaluating endogenous protein expression levels. Several CRISPR/Cas9 KO vectors were used to establish the effect of the targeted protein loss on the recruitment of EGFP or mCherry-fused DNA repair proteins to sites of laser-induced (micro-irradiation) DNA damage. All parental and modified cell lines were cultured in tissue culture incubators at 37°C, 10% CO_2_.

## Method Details

### Chemicals and Reagents

All chemicals and reagents used for these experiments are listed in the **Key Resources Table**. FK866 (NIMH #F-901; IUPAC name: (E)-[4-(1-Benzyoylpiperidin-4-yl)butly]-3-(pyridin-3-yl)acrylamide; CAS number: 201034-75-5) was obtained from the National Institute of Mental Health Chemical Synthesis and Drug Supply Program (Bethesda, MD). FK866 was dissolved in DMSO to prepare a stock solution at a concentration of 1 mM and stored at −80°C. Dihydronicotinamide Riboside (NRH; 1-[(2R,3R,4S,5R)-3,4-Dihydroxy-5-(hydroxymethyl)tetrahydrofuran-2-yl]-4H-pyridine-3-carboxamide) was prepared as described (Giroud-Gerbetant et al., 2019). NRH was dissolved in distilled H_2_O to prepare a stock solution concentration of 100 mM and stored at −80°C.

### Plasmid and vector development

Plasmids and lentiviral vectors developed previously or those newly generated for this study, either obtained commercially or from colleagues, are all cited in the **Key Resources Table** above. Lentiviral vectors were prepared by VectorBuilder Inc. unless specifically stated below. pLV-EGFP-Polβ-hygro-PAMmut (Sobol lab stock 1718), pLV-EGFP-Polβ(K72A)-hygro-PAMmut (Sobol lab stock 1719), and pLV-EGFP-Polβ(D256A)-hygro-PAMmut (Sobol lab stock 1720) were created by mutating nucleotide G24 in POLB (located in the sequence corresponding to exon 1) to G24A with the Quickchange II XL site-directed mutagenesis kit and the primers listed in the **Key Resources Table** to generate PAM mutants resistant CRISPR/Cas9 cleavage by POLBKO-gRNA1. Positive clones were selected and plasmids were extracted with the QIAprep Spin Miniprep Kit (Qiagen). Modifications were verified by Sanger sequencing (Eurofins Genomics).

The generation of the pUC19-POLBHR-EGFP insert used for endogenous tagging of the N-terminus of POLB in A549 cells began with modification of commercially available pUC19 plasmid to remove a SapI restriction site using the QuikChange II XL kit and PCR primers (see **Key Resource Table** - Sobol lab stock primers 1433 & 1434). Following clonal selection, the resulting plasmid was then modified to insert a ∼1.7kb high-fidelity PCR-amplified homology region fragment of POLB (including part of exon 1) generated using A549 genomic DNA as the template (Sobol lab stock primers 1427 & 1428). Following clonal selection, the resulting plasmid was modified to remove the PAM site in POLBHR used by the targeting CRISPR gRNA. Following clonal selection, the plasmid was modified via site-directed mutagenesis to add a SapI site located at the transcription start site of POLB (Sobol lab stock primers 1435 & 1436). An oligonucleotide was generated by high-fidelity PCR to contain SapI restriction fragments on the ends flanking the EGFP cDNA (Sobol lab stock primers 1437 & 1438). Lastly, the pUC19-POLBHR plasmid was modified to insert the SapI-EGFP DNA fragment via restriction digestion and ligation at the SapI site to produce a final pUC19 vector (pUC19-POLBHR-EGFP) with a ∼2.5kb insert containing ∼800bp upstream homology arm, EGFP in frame with the transcription start site of POLB, a PAM mutation in exon 1 to prevent secondary cleavage by CRISPR/Cas9, and ∼800bp downstream EGFP-POLB (see **Figure 2A**). The entire insert was sequenced via Sanger sequencing (Eurofins Genomics) to ensure proper generation.

Generation of constructs containing PAR binding domains (PBD) fused to EGFP began with a commercially purchased lentiviral backbone vector (VectorBuilder Inc.) containing a Gly-Ser linker and EGFP and MluI and BamHI restriction sites to enable in-frame cloning of the PBD (Sobol lab stock 1749). DNA fragments containing each of the PBD sequences were generated by GenScript USA, Inc (see **Supplemental Table S1**). PBD cDNA was amplified by high-fidelity PCR (Sobol lab stock primers 1485 & 1486). Each PBD-containing DNA fragment was ligated into the restriction-digested backbone vector and clonally selected. Insertion of the fragment was verified by Sanger sequencing (Eurofins Genomics) (see **Supplemental Table S2**).

### Lentivirus production and cell transduction

Lentiviral particles were generated by co-transfection of 4 plasmids into 293-FT cells using TransIT-X2 Transfection reagent: the packaging vectors pMD2.g(VSVG), pVSV-REV and pMDLg/pRRE together with the appropriate shuttle vectors, as listed in the **Key Resources Table** above. Forty-eight hours after transfection, lentivirus-containing supernatant was collected and passed through 0.45 mM filters to isolate the viral particles as described previously (Fang et al., 2014; Fouquerel et al., 2014).

Lentiviral transduction was performed as follows: cells (1-2x10^5^) were seeded into 6-well plates. 24 hrs later, lentiviral particles were mixed with polybrene (2µg/ml) and added to the cells. Cells were incubated at 32°C overnight and then medium with lentiviral particles was removed and replaced with fresh medium. When cells were created to form stable cell lines, cells were cultured for 48 hrs at 37°C before selection with antibiotics (puromycin or hygromycin) for 1-2 weeks. When cells were transduced a second time to create a cell expressing a fluorescently-tagged fusion protein in addition to harboring a KO, selection for the first stable cell line was completed and verified prior to initiation of the second transduction. When cells were created for transient expression experiments, cells were cultured for at least 96 hrs at 37°C before experimental analysis. All stable cell lines developed and used in this study are listed in the **Key Resource Table**.

### Development of Cas9 expressing and knockout (KO) cells

We developed U2OS and A549 cell lines with stable knockouts using the one vector CRISPR/Cas9 system (plentiCRISPR-v2; to deliver hSpcas9 and puromycin resistance). The plentiCRISPR-v2 vector was obtained from Addgene (plasmid #52961). The plentiCRISPR-v2 vectors containing control gRNA, POLB-KO-g2, XRCC1-KO-g1, XRCC1-KO-g2 and PARP1-KO-g1 were gifts from Wim Vermeulen (Erasmus MC, The Netherlands) (Slyskova et al., 2018). To generate plentiCRISPR-v2 containing gRNAs for POLB (using gRNA1), TARG, or SIRT6, we designed the guide RNA (gRNA) using the ChopChop software package (http://chopchop.cbu.uib.no). The resulting gRNAs (and the oligonucleotides used to generate the vectors) are listed in the **Key Resources Table**. The plasmids were used to generate lentivirus to express Cas9+Control-gRNA or Cas9+KO-gRNA, and U2OS or A549 cells were transduced with lentivirus as indicated above. Cells were maintained in media containing puromycin (1μg/ml) for 16 days, plated to generate single cell derived colonies, and validated by sequencing and protein immunoblot to confirm the knockout. Details of the technique have been described by us previously (Fang et al., 2019) and earlier by others (Sanjana et al., 2014).

### Generation of A549 cells expressing endogenous EGFP-tagged Polβ

Prior to use, the entire 2.5kb POLBHR-EGFP insert utilized for HR-dependent EGFP incorporation was enzymatically cleaved from the pUC19 backbone and gel purified to be used for CRISPR/Cas9 ribonucleotide protein insertion as below. To modify A549 cells to express EGFP-Polβ under control of the endogenous POLB promoter, the protocol provided by IDT was followed with minimal changes. First, tracrRNA (100 μM) and POLB gRNA1 (100 μM) were mixed in room temperature PBS, heated to 95°C, and cooled slowly to room temperature. Cas9 protein was diluted to a concentration of 1μM in room temperature PBS and then combined with the tracrRNA:POLBgRNA1 duplex to form the ribonucleoprotein complex in OptiMEM at room temperature for 5 minutes. The ribonucleoprotein complex was treated with RNAiMAX in OptiMEM for 20 minutes at room temperature to facilitate cellular delivery. During this time, the 2.5kb POLBHR-EGFP DNA fragment was treated with TransIT-X2 in Opti-MEM at room temperature for 15 minutes. Finally, A549 cells were trypsinized and reseeded to 5x10^5^ cells in a 6-well dish, followed by addition of the Cas9 ribonucleoprotein complex and the 2.5kb POLBHR-EGFP DNA fragment. Cells were incubated for 48 hours, at which time the cells were trypsinized and reseeded into glass-bottom 96-well dishes at single-cell density. Single cell clones were grown until visible colonies could be observed. Individual colonies were visualized for EGFP fluorescence using a Nikon A1rsi laser scanning confocal microscope to verify positive EGFP fluorescence. Positive colonies were trypsinized and grown for validation of CRISPR/Cas9 genomic modification of the POLB gene.

In addition to immunoblots and spectral immunofluorescence of EGFP-Polβ (methods below), Sanger sequencing was performed to verify correct insertion of EGFP onto the POLB gene. Genomic DNA from A549-POLBHR-EGFP cells was isolated, and the genomic region around the fragment insert site was PCR amplified using high-fidelity PCR and PCR-primers containing HindIII and EcoRI restriction fragments on the ends (Sobol lab stock primers 1427 & 1428). The PCR product was ligated into pUC19, transformed into STBL3 bacteria, and plated to obtain single colonies. Individual bacterial colonies were selected, and plasmids were isolated and sequenced completely across the POLBHR region to identify modifications in the POLBHR sequence. All three A549 POLB alleles were sequenced (see **Supplemental Figure 2**). Following validation, one cell clone was utilized for experimental investigation.

### Cell protein extract preparation

Cell protein extracts (whole cell lysates, WCL) were prepared from cells with different genetic modifications and/or treated with different drugs and for different times as indicated in the text. Cells were seeded into a 60-mm cell culture dish. After reaching 75-80% confluency, cells were washed twice with cold PBS, collected and lysed with an appropriate volume of 2x clear Laemmli buffer (2% SDS, 20% glycerol, 62.5mmol/l Tris-HCl pH6.8). Cell lysates were boiled for 10 min and quantified with the DC protein assay kit following the microplate protocol provided by the company (Bio-Rad).

### Immunoblot

Whole cell protein lysates (15-40μg protein) were loaded onto precast NuPAGE® Novex® 4-12% Bis-Tris gels, run 1hr at 120V. Gel electrophoresis separated proteins were transferred onto a PVDF membrane or nitrocellulose membrane using a Turboblotter (Bio-Rad). The membrane was first blocked with B-TBST (TBS buffer with 0.05% Tween-20 and supplemented with 5% blotting grade non-fat dry milk; Bio-Rad) for 1 hr at room temperature and subsequently blotted with the primary antibodies in B-TBST overnight at 4°C. The primary antibodies and their dilutions are listed in the **Key Resource Table**. After washing, membranes were incubated with secondary antibodies in B-TBST for 1 hr (room temperature). The following HRP conjugated secondary antibodies were used: Bio-Rad Goat anti-mouse-HRP conjugate and Bio-Rad anti-rabbit-HRP conjugate (see **Key Resource Table**). After washing, the membrane was illuminated with a chemiluminescent substrate. Protein bands were imaged using a Bio-Rad Chemi-Doc MP imaging system.

### MIDAS

**MIDAS** (for **M**odular **I**rradiation, **D**etection, and **A**nalysis **S**ystem) is a flexible, user-friendly and integrated software platform for start-to-finish performance and statistical analysis of micro-irradiation experiments. It is Modular in that each component offers the user multiple complementary approaches that may be freely combined. The Irradiation component is currently implemented as a macro written for NIS-Elements, which provides a graphical user interface (GUI) for choosing settings for laser wavelength, power, irradiation pattern(s), and image acquisition. Irradiation patterns and image acquisition can be tailored for time-lapse video of live cells expressing fluorescently labeled molecules of interest, for fixative-based staining following irradiation, or for combined approaches. Time-lapse irradiation experiments can be performed in parallel, with multiple cells in a single field being irradiated and then imaged collectively, or in series, where a selected group of cells are stimulated and imaged in sequence. Both approaches irradiate cells sequentially - the parallel module measures the duration of each irradiation event and applies a per-cell timing offset to accurately measure time post-irradiation when imaged. For fixative-based staining, a custom image registration algorithm is used to precisely re-locate the image field following sample preparation. After irradiation and image acquisition, the Detection component guides the user through the semi-automated measurement process, allowing for user supervision and intervention if necessary, while easing workflow by automating repetitive tasks. This component is implemented as a script for FIJI written in the Jython language, and features modes for measuring serial video, parallel video, or individual stained images. After data have been measured and output to a summary file, the Analysis component is used for visualization and statistical analysis of multiple datasets. This component is written in Python, using the Matplotlib and Numpy libraries for data processing. Data can be viewed at single-cell resolution or as averaged populations, with multiple normalization options to foreground different aspects of the data: normalized to a reference to minimize cell-to-cell variability, normalized to per-cell maximum intensity to emphasize differences in timing, and normalized to a pre-irradiation image to emphasize differences in intensity. To quantitatively assess features of recruitment data, three measurements are made for each intensity trace: time to peak recruitment intensity, half-life of recruitment, and relative peak intensity. Time to peak is defined as the time at which recruitment intensity crosses a threshold set to the 95% confidence interval of the maximum intensity for that cell. Half-life of recruitment is defined as the time post-peak at which the intensity crosses a lower threshold set to the same confidence interval of 50% of the maximum intensity. Relative peak intensity is defined as the ratio of maximum intensity per cell to the pre-irradiation intensity of that cell. Data are then analyzed statistically, and standard errors of the mean are reported. For record keeping and downstream analysis/visualization, an Excel format spreadsheet is generated, including all raw data, normalizations, measured features, statistical analyses, and experimental settings. To ensure data integrity and record keeping, although data may be excluded from analyses by the user, these data are still included in the summary spreadsheet, although plainly marked as excluded from analysis. MIDAS was developed and implemented at the USA Mitchell Cancer Institute (J.F.A.), with invaluable conceptual input from Dr. Natalie Gassman.

### Laser micro-irradiation

For laser micro-irradiation, 5x10^4^ cells were seeded into a 4-chamber glass bottom vessel (Thermo Fisher Scientific, #155382). 24 hours later, laser micro-irradiation was performed using a Nikon A1r confocal microscope. Live cells were imaged with a Nikon A1rsi laser scanning confocal microscope equipped with 6 visible wavelength lasers (405, 441, 514, 561, 647nm, Coherent), customized to add a UVA 355nm laser (PicoQuant) controlled by a Brueker XY Galvanometer, and equipped with a live-cell incubation chamber (Tokai Hit) maintained at 5% CO_2_ and 37°C, using a 40x oil-immersion objective. For each experiment, a field was selected, and cells were irradiated using a custom micro-irradiation script implemented in NIS-Elements. A 355nm laser or a 405nm laser (as indicated) was used for micro-irradiation, with stimulation times varying from 1s-2.5s per site. Time lapse images were collected every 15 seconds during a 10 minute interval. Images of focal recruitment were quantified using MIDAS to both detect focal recruitment and statistically analyze image results. Forty individual cells (2 sets of 10 cells were performed on 2 separate days) were analyzed and used to generate recruitment profiles and kinetic parameters.

### Immunofluorescence confocal microscopy

For poly(ADP-ribose) (PAR) analysis by immunocytochemistry, 5x10^4^ cells were seeded into a 4-chamber glass bottom vessel (Thermo Fisher Scientific, #155382). Cells were laser micro-irradiated at 405nm then subsequently fixed with 4% PFA for 10 min and briefly permeabilized with 0.1% TritonX-100 solution in PBS. Cells were rinsed with PBS and blocked in blocking buffer (10% normal goat serum in PBS) for 30 min and subsequently incubated with the PAR primary antibody for 1 to 2 hrs at 37°C, followed by three PBS washes and an incubation with fluorescent goat anti-mouse secondary antibody (see **Key Resources Table** for primary and antibodies used, with dilutions for each). Nuclei were stained with NucBlue dye. Fixed cells were imaged with a Nikon A1rsi laser scanning confocal microscope, using a 40x oil-immersion objective.

### Spectral Imaging of A549-POLBHR-EGFP cells

For spectral separation of low intensity EGFP visualization, spectral images were obtained with a Nikon A1rsi laser scanning confocal microscope using 488 nm laser for excitation, and collecting 14 bands from 500.2nm to 638.4nm, with a spectral gating resolution set to 10nm. Spectra were collected from parental and A549/EGFP-Polβ cells to provide spectra for autofluorescence and EGFP, respectively. Spectral unmixing was performed in NIS-Elements.

### NAD^+^/NADH Analysis

The level of NAD^+^ and NADH in cells was measured using the Enzychrome NAD^+^/NADH colorimetric assay kit (BioAssay Systems), following the supplier-provided protocols with minimal changes, as we have described previously (Wilk et al., 2020). Cells were seeded in a 6-well plate at a density of 2x10^5^ cells per well for NAD^+^ measurements and 3x10^5^ cells per well for NAD^+^ pool measurements (NAD^+^ plus NADH). 24 hrs later, cells were treated with the NRH (100 μM) for 0, 0.5, 1, 2, 4, 8 and 24 hours or with FK866 (50 nM) for 24 hours. Following treatment, cells were harvested and a suspension of 2x10^5^ cells was divided in half for measuring NAD^+^ and NADH, respectively, or a suspension of 1x10^5^ cells was used for the NAD^+^ measurement only. Cell pellets were homogenized using plastic pestles and the extraction of NAD^+^ and NADH was performed in the provided lysis buffers. Extracts were heated at 60°C for 5 min and neutralized with the provided buffers. Samples were spun down and the supernatant was immediately used for measurements of NAD^+^/NADH content using a Microplate Reader (BioTek) at 565 nm.

### Statistical analysis

Averages and standard error of the mean (SEM) were calculated from the means (on technical replicates) of multiple independent experiments (n = number of independent experiments as indicated in figure legends) unless stated otherwise. Student t-test and ANOVA was used to test for significant differences as appropriate, with results generally compared to controls and as indicated in the figure legends. P-values are indicated by asterisks with: *p<0.05, **p<0.01. Statistical analyses were performed using GraphPad PRISM except those explicitly determined in MIDAS.

## Notes

### Competing Interest Statement

The authors have declared no competing interest.

