## Supplement text and figures for "Temporal dynamics of base excision / single-strand break repair protein complex assembly and disassembly are modulated by the PARP1/NAD^+^/SIRT6 axis"

Affiliations
[1]Mitchell Cancer Institute, University of South Alabama, Mobile, AL 36604, USA
[2]Department of Pharmacology, College of Medicine, University of South Alabama, Mobile, AL 36604, USA

*Correspondence:
Robert W. Sobol, PhD; Mitchell Cancer Institute, University of South Alabama,
1660 Springhill Avenue, Mobile, AL 36604, USA.


**Figure S1: Laser-induced micro-irradiation of Polβ, XRCC1, and RealPAR**

A) Images of focal recruitment for 53BP1 (mApple-53BP1-trunc) in U2OS cells following 355nm and 405nm micro-irradiation. Images show a single cell either before (Pre-stim) or following 355nm or 405nm laser micro-irradiation at 0 min, 1 min, 5 min, 10 min or 20 minutes, post-stimulation. Recruitment of 53BP1 was observed as a bright red spot 5 minutes following 405nm stimulation, while no recruitment was observed following 355nm stimulation.

B) Recruitment of 53BP1 to sites of laser micro-irradiation (405nm) in U2OS cells, n ≥35. Error bars indicate standard error of the mean.

C) Images of focal recruitment for EGFP-Polβ, XRCC1-EGFP and RealPAR in U2OS cells following 355nm laser micro-irradiation. Focal recruitment can be observed as a bright green spot in each panel, with EGFP-Polβ and XRCC1-EGFP showing decreases in peak intensity over the time period indicated.

D) Recruitment of EGFP-Polβ to sites of laser micro-irradiation (355nm) in A549 cells, n ≥35. Error bars indicate standard error of the mean.

E) Recruitment of XRCC1-EGFP to sites of laser micro-irradiation (355nm) in A549 cells, n ≥35. Error bars indicate standard error of the mean.

F) Immunofluorescence of poly(ADP-ribose) (PAR) following laser micro-irradiation in EGFP-Polβ-expressing A549 cells. Cells were stimulated by laser micro-irradiation (355nm), fixed with formaldehyde, blocked and probed for PAR using a PAR monoclonal antibody (10H). Nuclei were stained with NucBlue, and cells were imaged. Merged images show colocalization of EGFP-Polβ (green) and PAR (red) at laser micro-irradiation sites.

G) Expression of PAR binding domain (PBD)-EGFP fusions in A549 cells. Each PBD was fused on the C-terminus with EGFP, each expressed in A549 cells following lentiviral transduction. Cells imaged after selection and stable cell line development with DAPI (nucleus) and phalloidin (actin filaments). Sequences for each PBD used can be found in **Supplement Table 1**.

H) Recruitment of RealPAR to sites of laser micro-irradiation (405nm) in A549 cells, n ≥35. RealPAR is recruited to laser micro-irradiation induced sites of DNA damage while the negative control (Y107A PAR binding deficient mutant) failed to recruit. Error bars indicate standard error of the mean.

**Figure S2: Over-expression of EGFP-Polβ displays similar recruitment kinetics as endogenously tagged EGFP-Polβ**

A) Full immunoblot images following analysis for Polβ, XRCC1, and PCNA (from **Figure 2C**) from lysates of A549 cells (C) and targeted A549 cells with endogenously expressed EGFP-Polβ (T). Molecular weight markers (kDa) are labeled on the left side. Arrows indicate the bands representing Polβ, EGFP-Polβ, XRCC1 or PCNA, as indicated.

B) Spectral imaging of A549 cells endogenously expressing EGFP-Polβ. From left to right: 1) a confocal image of A549 cells endogenously expressing EGFP-Polβ was obtained showing some perinuclear marks that were consistent with autofluorescence due to the low expression level of EGFP-Polβ; 2) a spectral image using all 14 channels were collected, with spectra from A549 parental and EGFP-Polβ-overexpressing A549 cells to provide spectra for autofluorescence and EGFP, respectively; 3) unmixed image displaying EGFP-Polβ in green and autofluorescence in red; 4) the unmixed image of EGFP-Polβ alone. Images show laser-induced micro-irradiation induced EGFP-Polβ recruitment as a bright green spot in multiple cells.

**Figure S3: Loss of Polβ enzymatic activity does not alter its recruitment kinetics**

A) Immunoblot of Polβ and actin of whole cell protein lysates prepared from A549/Cas9 control (C) and A549/POLB-KO (KO) cells. Molecular weight markers (kDa) are labeled on the left side.

B) Recruitment of EGFP-Polβ and the dRP lyase mutant, EGFP-Polβ(K72A), expressed in A549/POLB-KO cells, n $\geq$35. No change in recruitment was observed when endogenous POLB was knocked out. Error bars indicate standard error of the mean.

C) Time to peak recruitment intensity of EGFP-Polβ and EGFP-Polβ(K72A), following laser micro-irradiation. No significant difference was observed (Student's t-test). Error bars indicate standard error of the mean.

D) Half-life of recruitment of EGFP-Polβ and EGFP-Polβ(K72A), following laser micro-irradiation. No significant difference was observed (Student's t-test). Error bars indicate standard error of the mean.

**Figure S4: Polβ's recruitment is dependent on XRCC1, while Polβ enables XRCC1 complex dissociation**

Full gel immunoblot of CRISPR/Cas9-mediated XRCC1-KO from lysates of U2OS cells following expression of Cas9 or Cas9 plus the XRCC1-speific gRNA to exon2 or exon 3, as indicated. Expression of PCNA is shown as a loading control. Molecular weight markers (kDa) are labeled on the left side.

**Figure S5: Polβ and XRCC1 complex dynamics are dependent on PAR formation and degradation**

A) Immunoblot of TARG/OARD1 and PCNA from lysates of A549 cells following expression of Cas9 or Cas9 plus the TARG-specific gRNA, showing KO clones 1.2, 1.3 and 1.6.

B) Recruitment of EGFP-Polβ in A549/Cas9, A549/TARG-KO(1.3), and A549/TARG-KO(1.6) cells following 355nm micro-irradiation, $n \geq 35$. No change in recruitment was observed. Error bars indicate standard error of the mean.

**Figure S6: Polβ and XRCC1 complex dynamics are regulated by $NAD^+$ bioavailability**

A) Recruitment of EGFP-Polβ in U2OS cells, n ≥35, in media supplemented with either normal, non-heat inactivated FBS (black) or heat inactivated FBS. Following 355nm micro-irradiation, no change was observed. Error bars indicate standard error of the mean.

B) Recruitment of XRCC1-EGFP in U2OS cells, n ≥35, in media supplemented with either normal, non-heat inactivated FBS (black) or heat inactivated FBS. Following 355nm micro-irradiation, no change was observed. Error bars indicate standard error of the mean.

C) Recruitment of RealPAR in U2OS cells, n ≥35, in media supplemented with either normal, non-heat inactivated FBS (black) or heat inactivated FBS. Following 355nm micro-irradiation, no change was observed. Error bars indicate standard error of the mean.

D) NADH concentration in U2OS cells (left) and A549 cells (right), following NRH (100μM) treatment. No change in NADH was observed up to 8 hours following NRH treatment in either cell line.

E) Recruitment of EGFP-Polβ in A549 cells, n ≥35, following treatment with NRH (4 hrs, 100μM) or FK866 (24 hrs, 100μM). No change was observed following NRH treatment as compared to controls, while FK866 attenuated EGFP-Polβ recruitment to sites of 355nm micro-irradiation. Error bars indicate standard error of the mean.

F) Recruitment of XRCC1-EGFP in A549 cells, n ≥35, following treatment with NRH (4 hrs, 100μM) or FK866 (24 hrs, 100μM). No change was observed following NRH treatment as compared to controls, while FK866 attenuated XRCC1-EGFP recruitment to sites of 355nm micro-irradiation. Error bars indicate standard error of the mean.

G) Recruitment of RealPAR in A549 cells, n ≥35, following treatment with NRH (4 hrs, 100μM) or FK866 (24 hrs, 100μM). No change was observed following NRH treatment as compared to controls, while FK866 attenuated RealPAR recruitment to sites of 355nm micro-irradiation. Error bars indicate standard error of the mean.

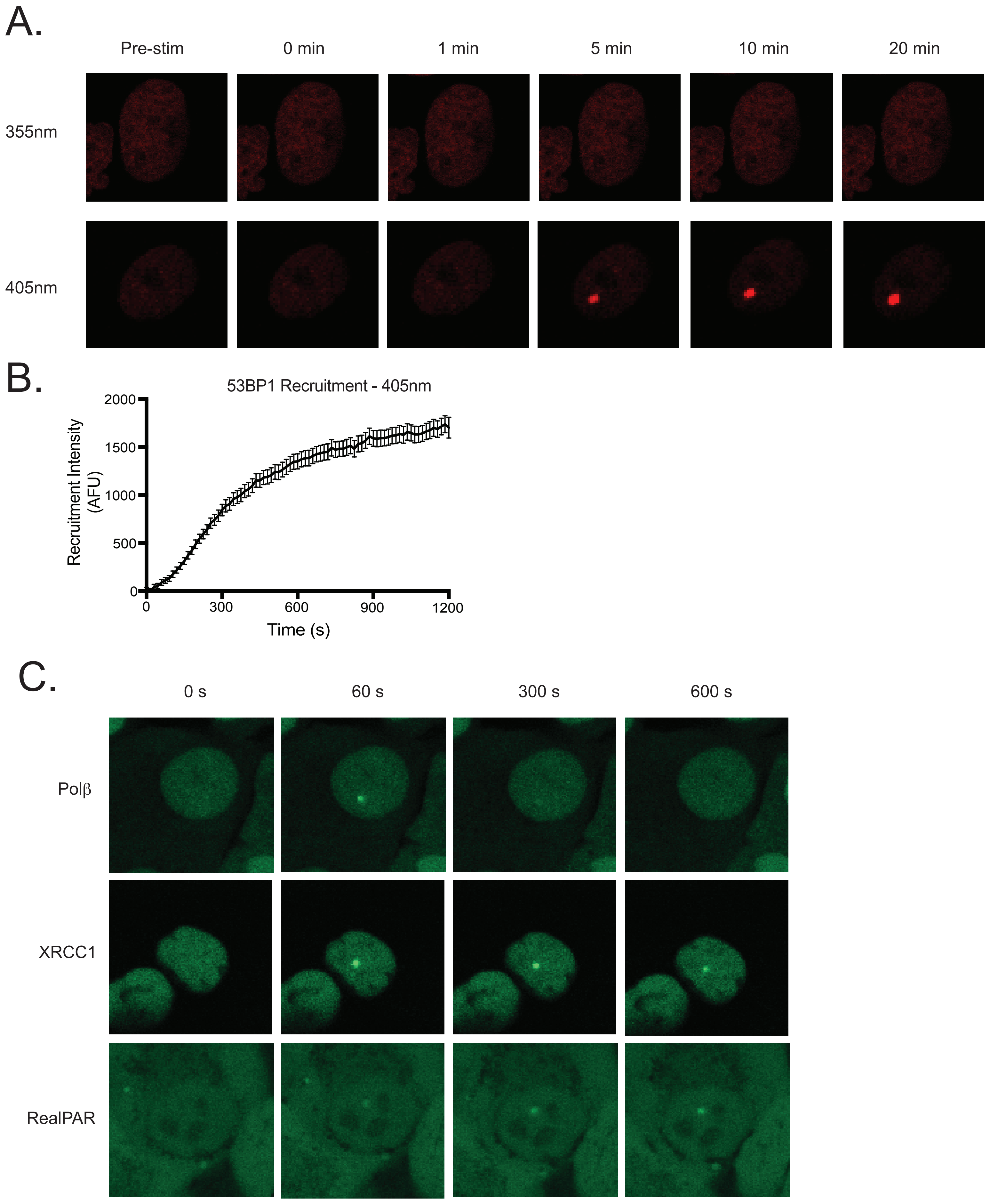
A.
Pre-stim
0 min
1 min
5 min
10 min
20 min
355nm
405nm
B.
53BP1 Recruitment - 405nm
Recruitment Intensity (AFU)
2000
1500
1000
500
0
0
300
600
900
1200
Time (s)
C.
0 s
60 s
300 s
600 s
Polβ
XRCC1
RealPAR

D.

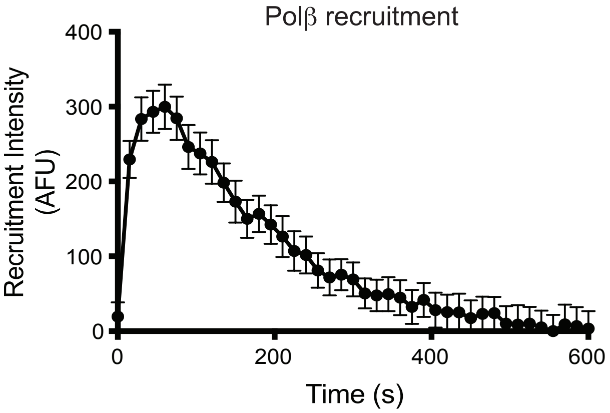


E.

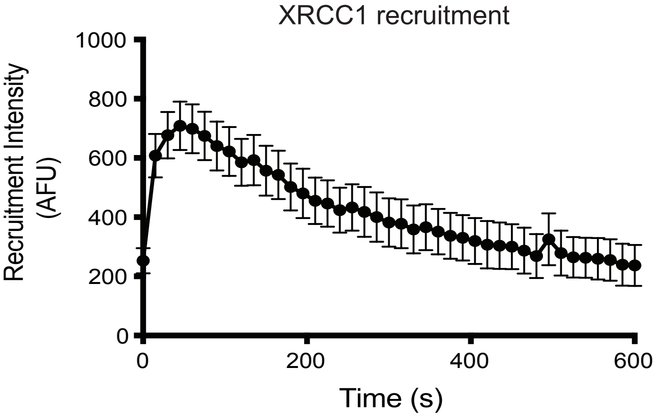


F.

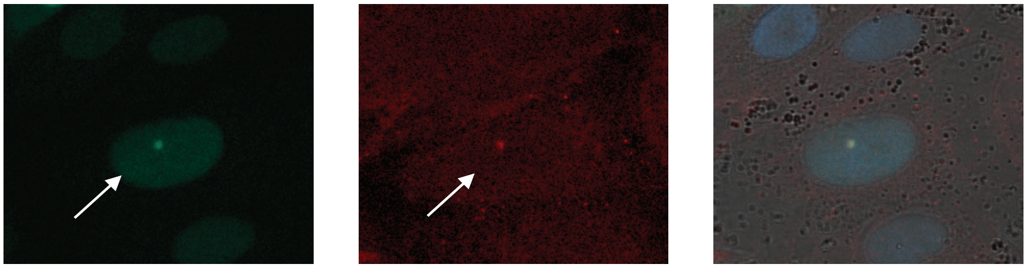

EGFP-Polβ

anti-PAR (10H)

Merged with NucBlue

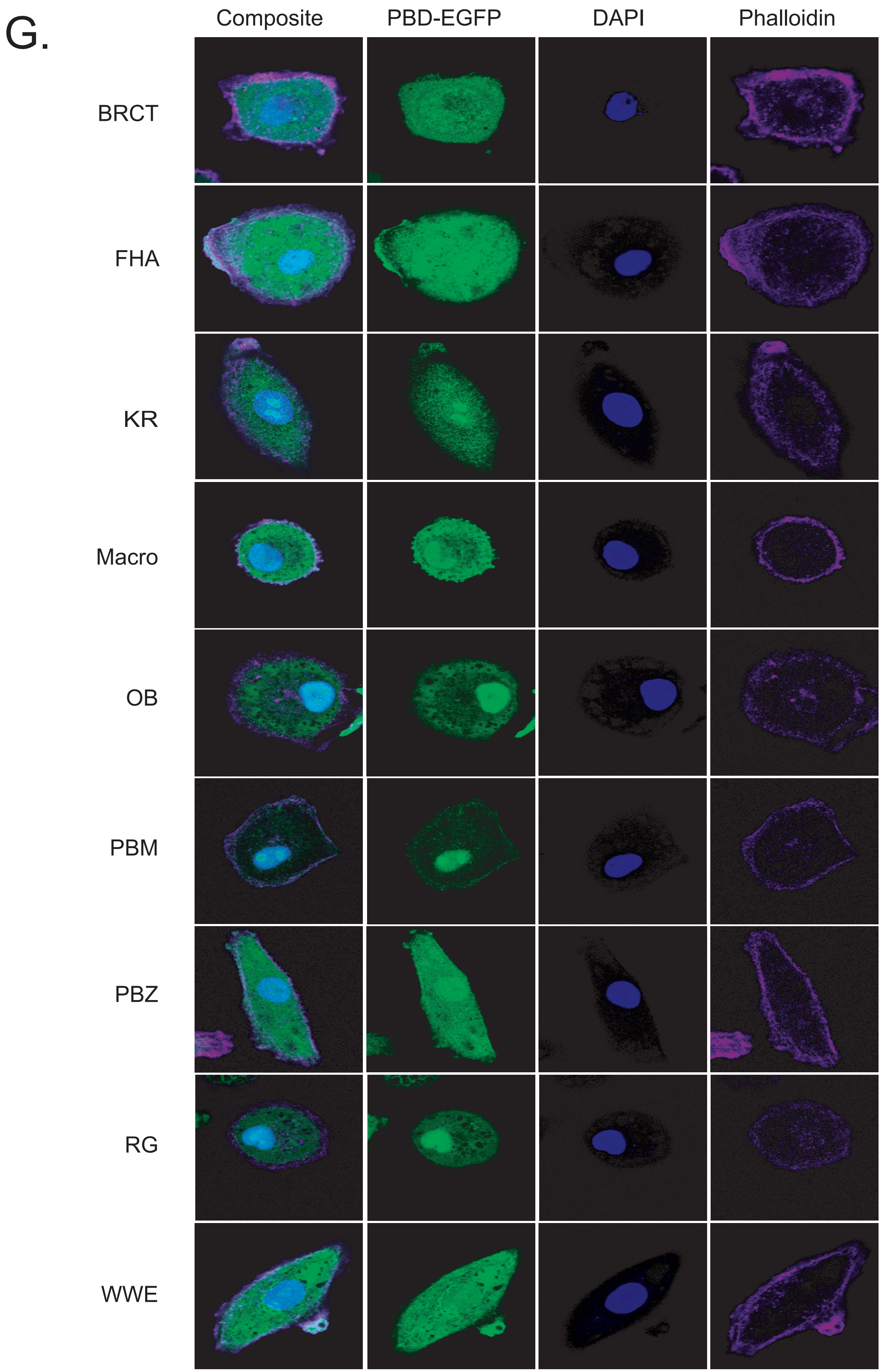
G.
Composite
PBD-EGFP
DAPI
Phalloidin
BRCT
FHA
KR
Macro
OB
PBM
PBZ
RG
WWE

H.

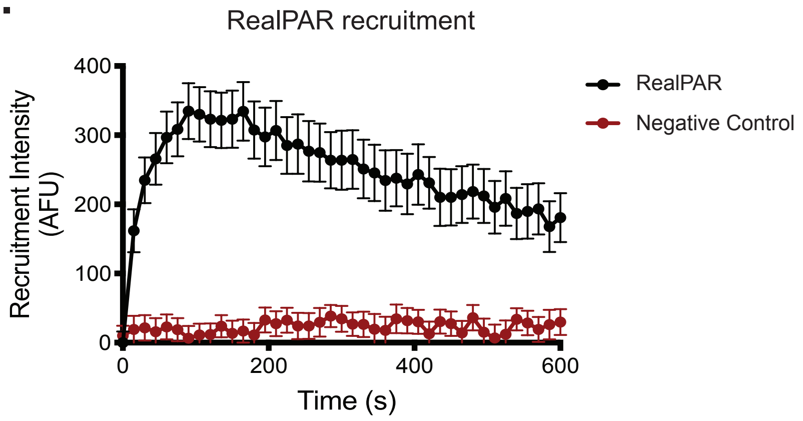
RealPAR recruitment
400
300
200
100
0
Recruitment Intensity (AFU)
0
200
400
600
Time (s)
RealPAR
Negative Control

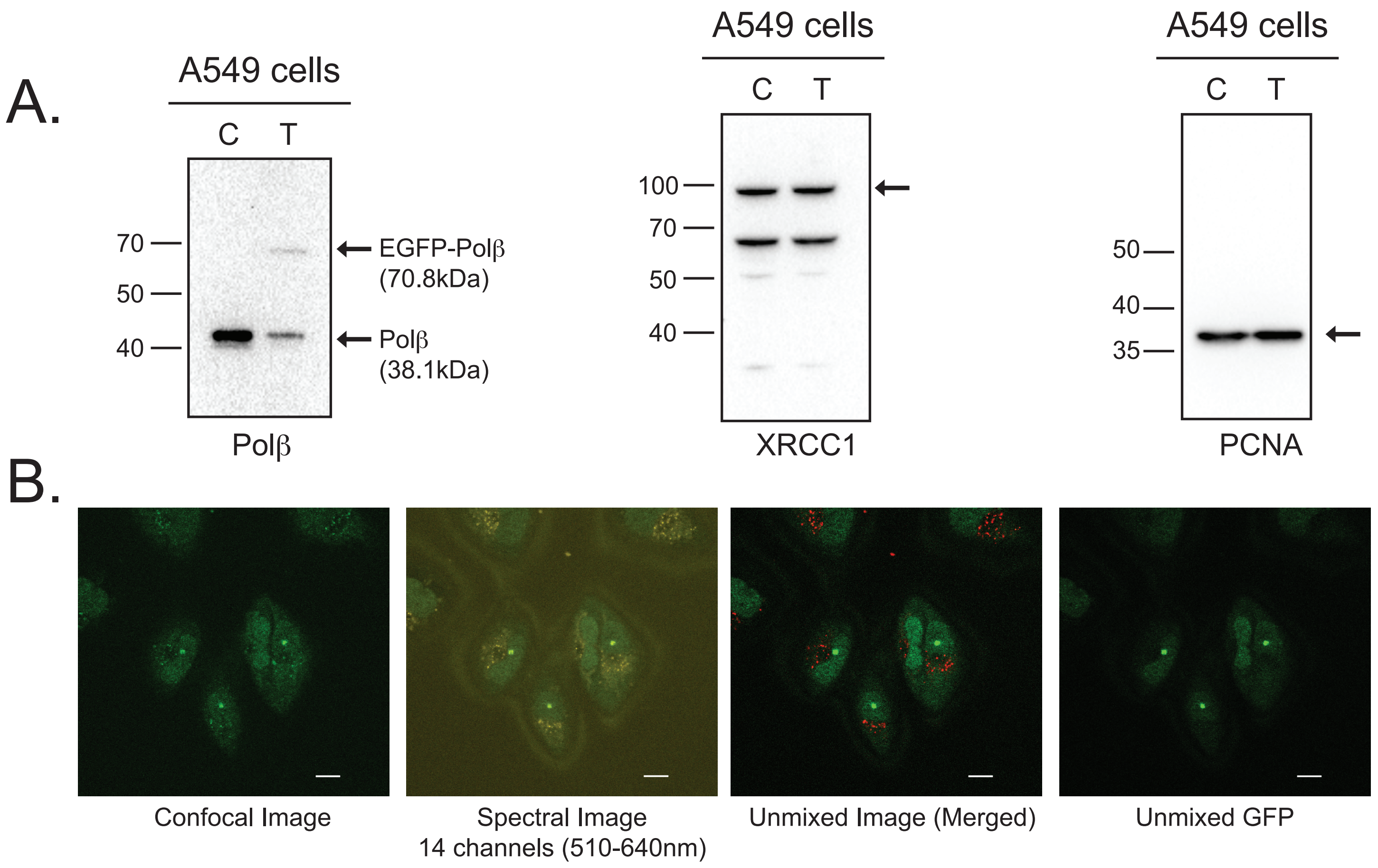
A.
A549 cells
C T
70
50
40
EGFP-Polβ (70.8kDa)
Polβ (38.1kDa)
Polβ
A549 cells
C T
100
70
50
40
XRCC1
A549 cells
C T
50
40
35
PCNA
B.
Confocal Image
Spectral Image 14 channels (510-640nm)
Unmixed Image (Merged)
Unmixed GFP

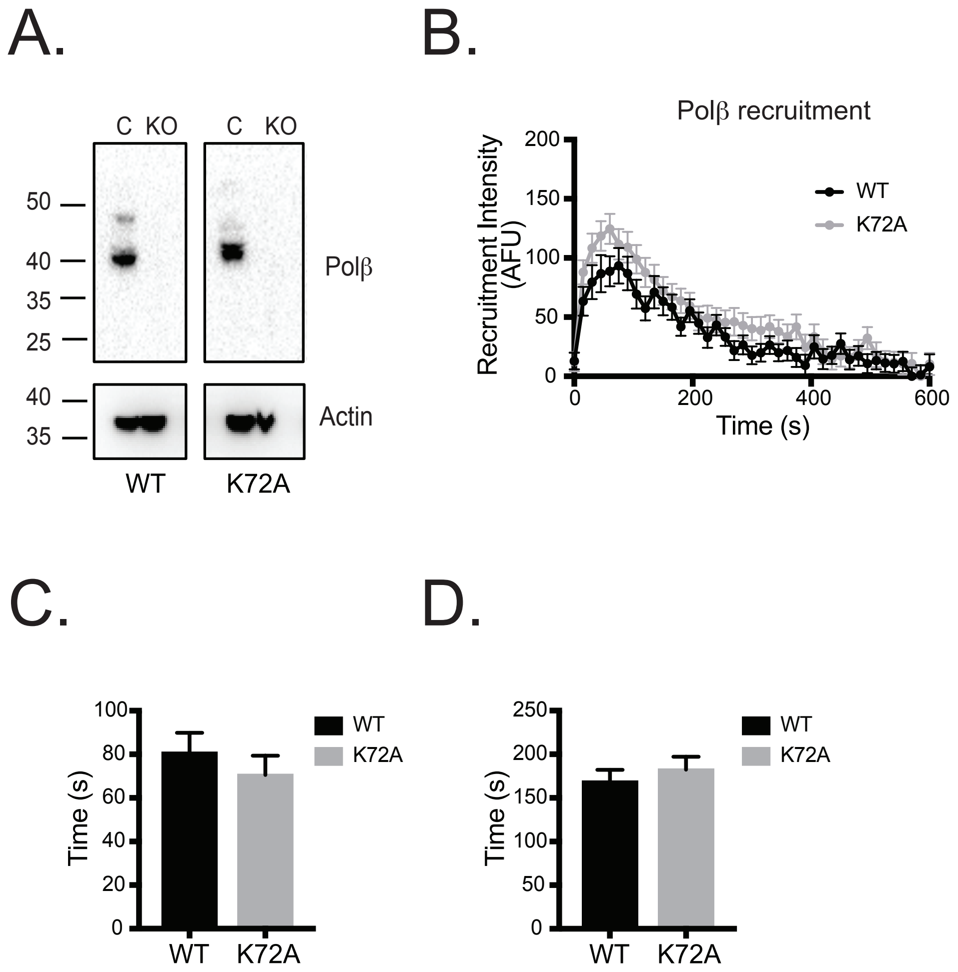
A.
C KO
C KO
50
40
35
25
Polβ
40
35
Actin
WT
K72A
B.
Polβ recruitment
Recruitment Intensity (AFU)
200
150
100
50
0
WT
K72A
0
200
400
600
Time (s)
C.
Time (s)
100
80
60
40
20
0
WT
K72A
WT
K72A
D.
Time (s)
250
200
150
100
50
0
WT
K72A
WT
K72A

A.

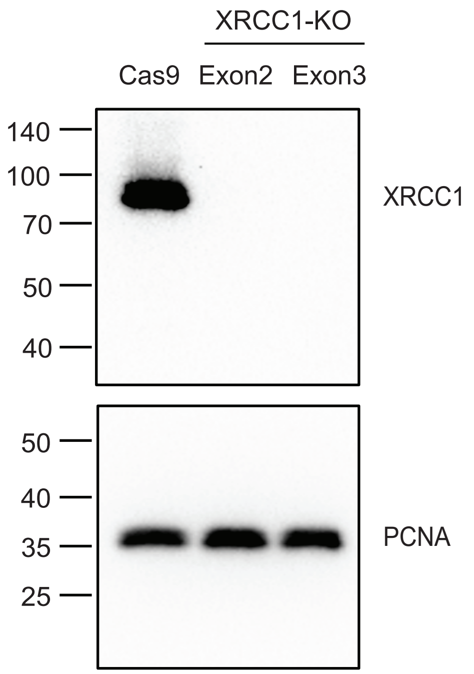
XRCC1-KO
Cas9
Exon2
Exon3
140
100
70
50
40
XRCC1
50
40
35
25
PCNA

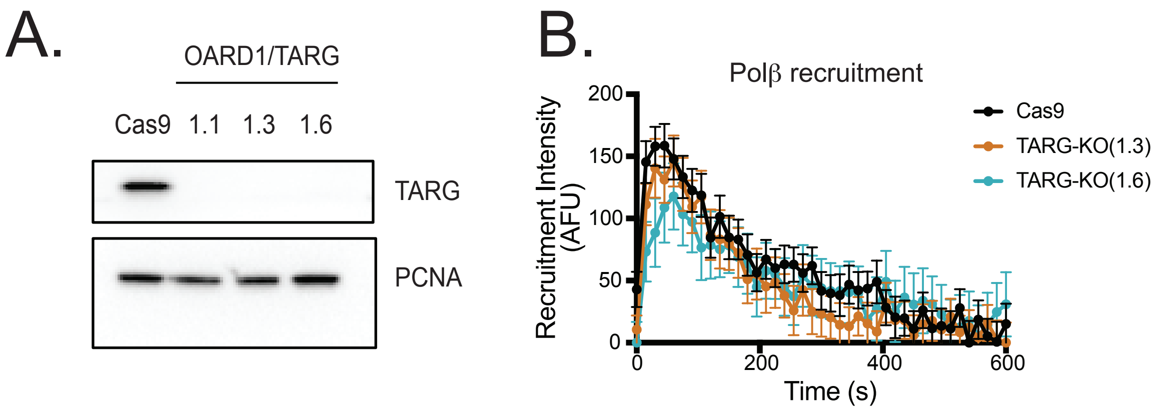
A.
OARD1/TARG
Cas9
1.1
1.3
1.6
TARG
PCNA
B.
Polβ recruitment
Recruitment Intensity (AFU)
200
150
100
50
0
0
200
400
600
Time (s)
Cas9
TARG-KO(1.3)
TARG-KO(1.6)

Koczor et al.

Supplement: Figure S6

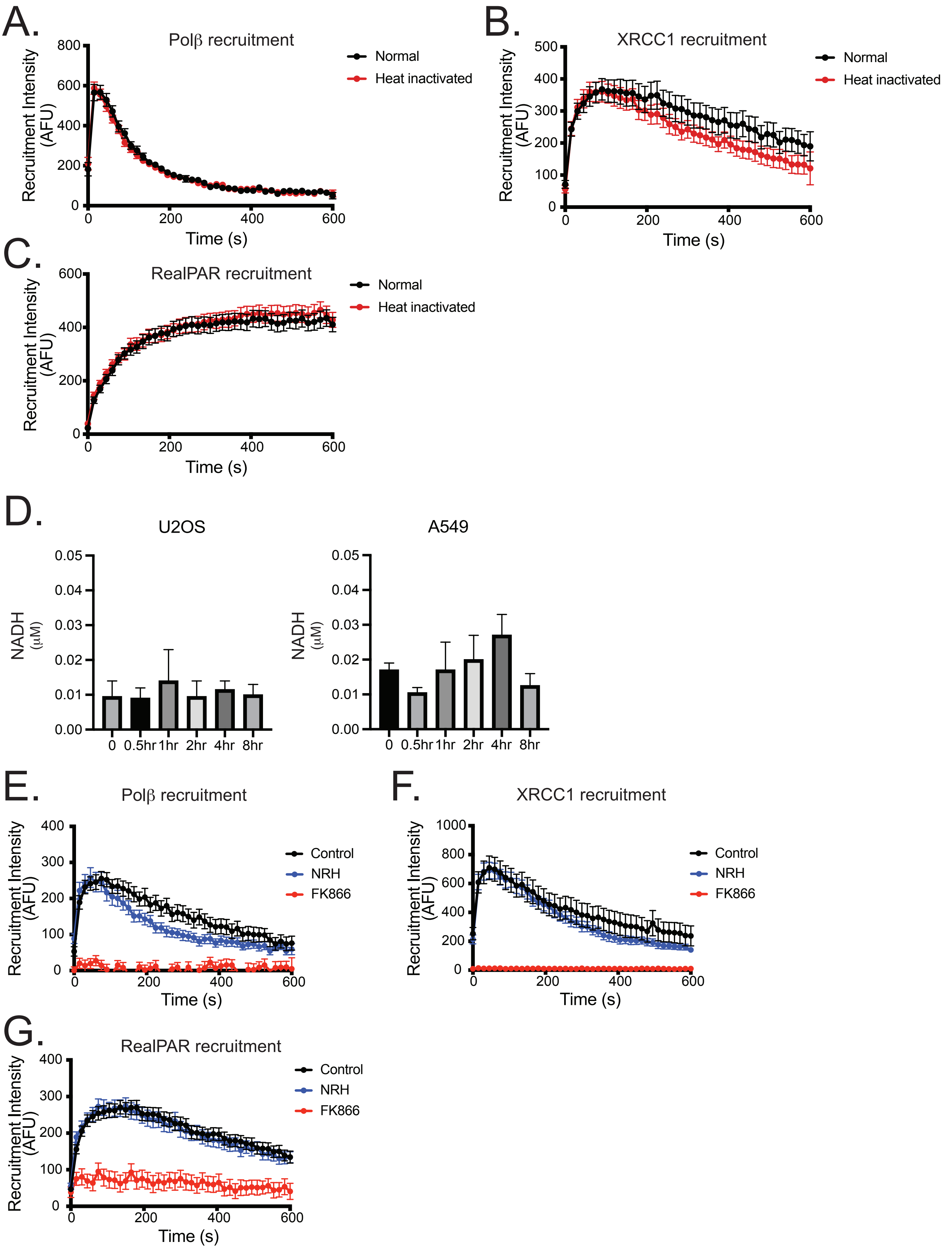

**Table S1: PAR binding domains (PBD)**

| PBD | Origin of PBD Sequence | 5' Overhang | MluI Site | Kozak | Start Codon | PBD Sequence | Linker | BamHI Site | 3' Overhang | Complete Sequence | Length (bp) |
|---|---|---|---|---|---|---|---|---|---|---|---|
| BRCT | XRCC1 | GTGGTT | ACGCGT | GCCACC | ATG | GTAGTGGTGCTGAGTGGCTTCCAGAACCCCTTC<br>CGCTCCGAGCTGCGAGATAAGGCCCTAGAGCTT<br>GGGGCCAAGTATCGGCCAGACTGGACCCGGGA<br>CAGCACGCACCTCATCTGTGCCTTTGCCAACACC<br>CCCAAGTACAGCCAGGTCCTAGGC | TCTGGTGGCGGAGGCTCGGGCGGAGGT | GGATCC | TCCTTC | GTGGTTACGCGTGCCACCATGGTAGTGGTGCTGAGTG<br>GCTTCCAGAACCCCTTCCGCTCCGAGCTGCGAGATAAG<br>GCCCTAGAGCTTGGGGCCAAGTATCGGCCAGACTGGA<br>CCCGGGACAGCACGCACCTCATCTGTGCCTTTGCCAAC<br>ACCCCCAAGTACAGCCAGGTCCTAGGCTCTGGTGGCG<br>GAGGCTCGGGCGGAGGTGGATCCTCCTTC | 216 |
| FHA | APTX | GTGGTT | ACGCGT | GCCACC | ATG | GCAGTTGTGATTGGGCGTGGCCCAGAGACCAA<br>GATCACTGATAAGAAATGTTCTCGACAGCAAGT<br>ACAGTTGAAAGCAGAG | TCTGGTGGCGGAGGCTCGGGCGGAGGT | GGATCC | TCCTTC | GTGGTTACGCGTGCCACCATGGCAGTTGTGATTGGGC<br>GTGGCCCAGAGACCAAGATCACTGATAAGAAATGTTC<br>TCGACAGCAAGTACAGTTGAAAGCAGAGTCTGGTGGC<br>GGAGGCTCGGGCGGAGGTGGATCCTCCTTC | 141 |
| KR | dMI-2 | GTGGTT | ACGCGT | GCCACC | ATG | CGCAGGAAGAAGAAGGGCAAGAAGCGCAAGA<br>CCAGGAAGGGCGAAGAGAAGGGTCGCAAGAA<br>GAAGAAGCGCAAGAAG | TCTGGTGGCGGAGGCTCGGGCGGAGGT | GGATCC | TCCTTC | GTGGTTACGCGTGCCACCATGCGCAGGAAGAAGAAG<br>GGCAAGAAGCGCAAGACCAGGAAGGGCGAAGAGAA<br>GGGTCGCAAGAAGAAGAAGCGCAAGAAGTCTGGTG<br>GCGGAGGCTCGGGCGGAGGTGGATCCTCCTTC | 138 |
| Macro | H2A1.1 | GTGGTT | ACGCGT | GCCACC | ATG | ACAGTCCTCTCCACCAAGAGCCTCTTCCTTGGCC<br>AGAAGTTGCAAGTTGTACAGGCTGACATTGCCT<br>CGATCGACAGTGATGCTGTCGTTCACCCGACAA<br>ACACTGACTTCTACATCGGTGGTGAAGTAGGAA<br>ACACGCTGGAGAAGAAAGGTGGCAAGGAGTT<br>TGTGGAAGCTGTCCTGGAACTCCGGAAAAAGA<br>ACGGGCCCTTGGAAGTAGCTGGAGCTGCTGTC<br>AGCGCAGGCCATGGCCTGCCTGCCAAGTTTGTG<br>ATCCACTGTAATAGTCCAGTTTGGGGTGCAGAC<br>AAGTGTGAAGAACTTCTGGAAAAGACAGTGAA<br>AAACTGCTTGGCCCTGGCTGATGATAAGAAGCT<br>GAAATCCATTGCATTTCCATCCATCGGCAGCGGC<br>AGGAACGGTTTTCCAAAGCAGACAGCAGCTCA<br>GCTGATTCTGAAGGCCATCTCCAGTTACTTCGTG<br>TCTACAATGTCCTCTTCCATCAAAACGGTGTACT<br>TCGTGCTTTTTGACAGCGAGAGTATAGGCATCT<br>ATGTGCAGGAAATGGCCAAGCTGGACGCCAAC | TCTGGTGGCGGAGGCTCGGGCGGAGGT | GGATCC | TCCTTC | GTGGTTACGCGTGCCACCATGACAGTCCTCTCCACCAA<br>GAGCCTCTTCCTTGGCCAGAAGTTGCAAGTTGTACAG<br>GCTGACATTGCCTCGATCGACAGTGATGCTGTCGTTCA<br>CCCGACAAACACTGACTTCTACATCGGTGGTGAAGTAG<br>GAAACACGCTGGAGAAGAAAGGTGGCAAGGAGTTTG<br>TGGAAGCTGTCCTGGAACTCCGGAAAAAGAACGGGCC<br>CTTGGAAGTAGCTGGAGCTGCTGTCAGCGCAGGCCAT<br>GGCCTGCCTGCCAAGTTTGTGATCCACTGTAATAGTCC<br>AGTTTGGGGTGCAGACAAGTGTGAAGAACTTCTGGA<br>AAAGACAGTGAAAAACTGCTTGGCCCTGGCTGATGAT<br>AAGAAGCTGAAATCCATTGCATTTCCATCCATCGGCAG<br>CGGCAGGAACGGTTTTCCAAAGCAGACAGCAGCTCAG<br>CTGATTCTGAAGGCCATCTCCAGTTACTTCGTGTCTACA<br>ATGTCCTCTTCCATCAAAACGGTGTACTTCGTGCTTTTT<br>GACAGCGAGAGTATAGGCATCTATGTGCAGGAAATG<br>GCCAAGCTGGACGCCAACTCTGGTGGCGGAGGCTCG<br>GGCGGAGGTGGATCCTCCTTC | 618 |
| OB | SSB1 | GTGGTT | ACGCGT | GCCACC | ATG | ACGACGGAGACCTTTGTGAAGGATATCAAGCCT<br>GGGCTCAAGAATCTGAACCTTATCTTCATTGTGC<br>TGGAGACAGGCCGAGTGACCAAGACAAAGGAC<br>GGGCATGAGGTTCGGACCTGCAAAGTGGCGG<br>ACAAAACAGGCAGCATCAATATCTCTGTCTGGG<br>ACGATGTTGGCAATCTGATCCAGCCTGGGGACA<br>TTATCCGGCTCACCAAAGGGTACGCTTCAGTTTT<br>CAAAGGTTGTCTGACACTATATACTGGCCGTGG<br>GGGTGATCTGCAGAAGATTGGAGAATTCTGTA<br>TGGTTTATTCTGAGGTTCCTAACTTCAGTGAG | TCTGGTGGCGGAGGCTCGGGCGGAGGT | GGATCC | TCCTTC | GTGGTTACGCGTGCCACCATGACGACGGAGACCTTTG<br>TGAAGGATATCAAGCCTGGGCTCAAGAATCTGAACCT<br>TATCTTCATTGTGCTGGAGACAGGCCGAGTGACCAAG<br>ACAAAGGACGGGCATGAGGTTCGGACCTGCAAAGTG<br>GCGGACAAAACAGGCAGCATCAATATCTCTGTCTGGG<br>ACGATGTTGGCAATCTGATCCAGCCTGGGGACATTATC<br>CGGCTCACCAAAGGGTACGCTTCAGTTTTCAAAGGTTG<br>TCTGACACTATATACTGGCCGTGGGGGTGATCTGCAG<br>AAGATTGGAGAATTCTGTATGGTTTATTCTGAGGTTCC<br>TAACTTCAGTGAGTCTGGTGGCGGAGGCTCGGGCGG<br>AGGTGGATCCTCCTTC | 387 |
| PBM | Concensus PBD<br>(Pleishke et al., 2000) | GTGGTT | ACGCGT | GCCACC | ATG | AAGAAGAAGAAGAAGAGGTTCAGCTTCAAGAA<br>GAGCTTCAAGCTGAGCGGCTTCAGCTTCAAGAA<br>GAACAAGAAG | TCTGGTGGCGGAGGCTCGGGCGGAGGT | GGATCC | TCCTTC | GTGGTTACGCGTGCCACCATGAAGAAGAAGAAGAAG<br>AGGTTCAGCTTCAAGAAGAGCTTCAAGCTGAGCGGCT<br>TCAGCTTCAAGAAGAACAAGAAGTCTGGTGGCGGAG<br>GCTCGGGCGGAGGTGGATCCTCCTTC | 135 |
| PBZ-C | CHFR | GTGGTT | ACGCGT | GCCACC | ATG | CGTCCTGACTGCTACTGGGGCCGTAACTGCCGC<br>ACTCAGGTGAAAGCTCACCACGCCATGAAATTC<br>AATCATATC | TCTGGTGGCGGAGGCTCGGGCGGAGGT | GGATCC | TCCTTC | GTGGTTACGCGTGCCACCATGCGTCCTGACTGCTACTG<br>GGGCCGTAACTGCCGCACTCAGGTGAAAGCTCACCAC<br>GCCATGAAATTCAATCATATCTCTGGTGGCGGAGGCTC<br>GGGCGGAGGTGGATCCTCCTTC | 135 |
| RG | MRE11 | GTGGTT | ACGCGT | GCCACC | ATG | GATGATAGCATCTCAGCAGCAACCAACAAAGGA<br>AGAGGCCGAGGAAGAGGTCGAAGAGGTGGA<br>AGAGGGCAGAATTCAGCATCGAGAGGAGGGT<br>CTCAA | TCTGGTGGCGGAGGCTCGGGCGGAGGT | GGATCC | TCCTTC | GTGGTTACGCGTGCCACCATGGATGATAGCATCTCAG<br>CAGCAACCAACAAAGGAAGAGGCCGAGGAAGAGGTC<br>GAAGAGGTGGAAGAGGGCAGAATTCAGCATCGAGA<br>GGAGGGTCTCAATCTGGTGGCGGAGGCTCGGGCGGA<br>GGTGGATCCTCCTTC | 159 |
| WWE | RNF146 | GTGGTT | ACGCGT | GCCACC | ATG | ATGGGAAATGGTGAATATGCATGGTATTATGA<br>AGGAAGAAATGGGTGGTGGCAGTACGATGAG<br>CGCACTAGTAGAGAGCTGGAAGATGCTTTTTCC<br>AAAGGTAAAAAGAACACTGAAATGTTAATTGCT<br>GGCTTTCTGTATGTCGCTGATCTTGAAAACATG<br>GTTCAATATAGGAGAAATGAACATGGACGTCG<br>CAGGAAGATTAAGCGAGATATAATAGATATAC<br>CAAAGAAGGGAGTAGCTGGACTTAGG | TCTGGTGGCGGAGGCTCGGGCGGAGGT | GGATCC | TCCTTC | GTGGTTACGCGTGCCACCATGATGGGAAATGGTGAAT<br>ATGCATGGTATTATGAAGGAAGAAATGGGTGGTGGC<br>AGTACGATGAGCGCACTAGTAGAGAGCTGGAAGATG<br>CTTTTTCCAAAGGTAAAAAGAACACTGAAATGTTAATT<br>GCTGGCTTTCTGTATGTCGCTGATCTTGAAAACATGGT<br>TCAATATAGGAGAAATGAACATGGACGTCGCAGGAA<br>GATTAAGCGAGATATAATAGATATACCAAAGAAGGG<br>AGTAGCTGGACTTAGGTCTGGTGGCGGAGGCTCGGG<br>CGGAGGTGGATCCTCCTTC | 312 |

**Table S2: Endogenously-tagged EGFP-POLB Allele Sequences**

Key: **Homology Regions** | EGFP | POLB Exon 1 | PAM Mutation | Possible Coding (18bp starting at ATG)

Allele 1: No modification

TTGTAAAATAAACTATTGCTGATGTGACATTTCCATTTCTCAAGCAGACTGTAAATTTCTCAGATGTAGGTCATCATGTTATAATTAGGGTTCCATTGCTCATGATCTAGATGAAACACTTCTCCCTTTCTTGTGGTCTCTT
CATTAATTTGGTCCCTACCGCAAAGGCAGGGGGATTAAAAATAAAATATCTTTTGACACTGGTTGGCAGGTGTACTAACACGTGTGCGGGAGGATTCCTGTTCTCGGCATGGTTCACGCCGGGCTCCGGCAGCGCC
CTAGTACGTAAGGGGCTGCACTGGAGTGTGGACTGCCCGGTATCCTTAGAGAAAGTTTTGACAGTGTGACGCCAGTGTCGCCTCTAACCACGTACAAAACGCGCCATGCATTGGGTAGCTATTACAGTGCACATTTT
TGTGCAGGAAGCTACCCCACCAGCGCAAGTATTTCAGAATCAAGATCGCACTCCCGTTTCCCCTTCTAGGGAAAGGATTCCAGATAAACACTGAGAAAACAGATTTAATATCTTAGGTCAAAACTCCAGGTCTTCCCA
TAGGAAGGCCCCTGGGCTGTCATTCTGAGGTGCCTATTTCCCCCGCGCCGTGTCCACGTCCACGAGTCCACGAACCCTCCGAGTTCTCTCTCCTCACCTGCCGAGGGCTCGCCCCTCTAGCCCCGCCCCGCCCCGC
CCAGGACGCGTGACGTCACAACAAGCGCCGTGCGCCCCGCCCCCCATCGGGGGCAACCATTGTTCCGCCGGTCGCGCCGGAGCTGGGTTGCTCCTGCTCCCGTCTCCAAGTCCTGGTACCTCCTTCAAGCTGGGA
GAGGGCTCTAGTCCCTGGTTCTGAACACTCTGGGGTTCTCGGGTGCAGGCCGCCATGAGCAAACGGAGGGCGCCGCAAGAGACTCTCAACGGGGGAATCACCGACATGCTCACAGGTTAGCACCGGGCCGGGC
CCCGCTGGCTTTCTTCTTTCCTTCCAGCCTCTTCCCCTGCCTTCCTTCTCTCCCACACCGACAGTCCAGTGGGTAGGGTAGGTTCCTTGCAGCGGGTCGTCTTCCGTGGGGATCTCCCTCCGGCGCCCCTGGCTGGTT
GTCAGTCCTGCCGGCTACACCTGGGCCATCGCTTGGGCTGCTTTTGGTCTGGCCCTTGGAGGAAACGGGTGGTCACTGTCTTTTAGAGGCTGCCATATCCCCTTCCAGAAAACAGTTCTCGTTTGTTTACTCGTGGTT
CTTGTTCACCCGAGCCTTCTGTTGCCTTTCAGAACTCGCAAACTTTGAGAAGAACGTGAGCCAAGCTATCCACAAGTACAATGCTTACAGGTGGGACAGTGCAGCATTCTCGGGTAGCATACGTTCTGGGATACCCT
GTTTAGTGTGGCAATTAACAGGACTGAGGGCCCAGTGGATATTTGGTCCATCTGCAAGAGCGGGAAAAAGCAAGAATCGAGGCTGGTACCTTACTATTTCTTGAAGAATGTGGGCAGTGCGTTATAGGATCAGTTC
AGATAAGTTCCCAATTCTGATTGCAGATAGCTGTAGCCTGATACGTATTTTGTCATCATCAAACATGAATTACTAGAAATGCGGTGTAATTTTAACCGTAATGAAAGACAGCGGATGAAGTCCATACTGTGAAATATAT
GTGGTTCAAGGAATCACATTTTTTCTTTTGGTTGGTCTAAAGTTGATCCTGTGGGATCCTGATCCTGATCCTTGTTCCTT

Allele 2: Full EGFP insertion

TTGTAAAATAAACTATTGCTGATGTGACATTTCCATTTCTCAAGCAGACTGTAAATTTCTCAGATGTAGGTCATCATGTTATAATTAGGGTTCCATTGCTCATGATCTAGATGAAACACTTCTCCCTTTCTTGTGGTCTCTT
CATTAATTTGGTCCCTACCGCAAAGGCAGGGGGATTAAAAATAAAATATCTTTTGACACTGGTTGGCAGGTGTACTAACACGTGTGCGGGAGGATTCCTGTTCTCGGCATGGTTCACGCCGGGCTCCGGCAGCGCC
CTAGTACGTAAGGGGCTGCACTGGAGTGTGGACTGCCCGGTATCCTTAGAGAAAGTTTTGACAGTGTGACGCCAGTGTCGCCTCTAACCACGTACAAAACGCGCCATGCATTGGGTAGCTATTACAGTGCACATTTT
TGTGCAGGAAGCTACCCCACCAGCGCAAGTATTTCAGAATCAAGATCGCACTCCCGTTTCCCCTTCTAGGGAAAGGATTCCAGATAAACACTGAGAAAACAGATTTAATATCTTAGGTCAAAACTCCAGGTCTTCCCA
TAGGAAGGCCCCTGGGCTGTCATTCTGAGGTGCCTATTTCCCCCGCGCCGTGTCCACGTCCACGAGTCCACGAACCCTCCGAGTTCTCTCTCCTCACCTGCCGAGGGCTCGCCCCTCTAGCCCCGCCCCGCCCCGC
CCAGGACGCGTGACGTCACAACAAGCGCCGTGCGCCCCGCCCCCCATCGGGGGCAACCATTGTTCCGCCGGTCGCGCCGGAGCTGGGTTGCTCCTGCTCCCGTCTCCAAGTCCTGGTACCTCCTTCAAGCTGGGA
GAGGGCTCTAGTCCCTGGTTCTGAACACTCTGGGGTTCTCGGGTGCAGGCCGCCATGGTGAGCAAGGGCGAGGAGCTGTTCACCGGGGTGGTGCCCATCCTGGTCGAGCTGGACGGCGACGTAAACGGCCACA
AGTTCAGCGTGTCCGGCGAGGGCGAGGGCGATGCCACCTACGGCAAGCTGACCCTGAAGTTCATCTGCACCACCGGCAAGCTGCCCGTGCCCTGGCCCACCCTCGTGACCACCCTGACCTACGGCGTGCAGTGC
TTCAGCCGCTACCCCGACCACATGAAGCAGCACGACTTCTTCAAGTCCGCCATGCCCGAAGGCTACGTCCAGGAGCGCACCATCTTCTTCAAGGACGACGGCAACTACAAGACCCGCGCCGAGGTGAAGTTCGAG
GGCGACACCCTGGTGAACCGCATCGAGCTGAAGGGCATCGACTTCAAGGAGGACGGCAACATCCTGGGGCACAAGCTGGAGTACAACTACAACAGCCACAACGTCTATATCATGGCCGACAAGCAGAAGAACGG
CATCAAGGTGAACTTCAAGATCCGCCACAACATCGAGGACGGCAGCGTGCAGCTCGCCGACCACTACCAGCAGAACACCCCCATCGGCGACGGCCCCGTGCTGCTGCCCGACAACCACTACCTGAGCACCCAGTC
CGCCCTGAGCAAAGACCCCAACGAGAAGCGCGATCACATGGTCCTGCTGGAGTTCGTGACCGCCGCCGGGATCACTCTCGGCATGGACGAGCTGTACAAGATGAGCAAACGGAAGGCGCCGCAAGAGACTCTC
AACGGGGGAATCACCGACATGCTCACAGGTTAGCACCGGGCCGGGCCCCGCTGGCTTTCTTCTTTCCTTCCAGCCTCTTCCCCTGCCTTCCTTCTCTCCCACACCGACAGTCCAGTGGGTAGGGTAGGTTCCTTGCA
GCGGGTCGTCTTCCGTGGGGATCTCCCTCCGGCGCCCCTGGCTGGTTGTCAGTCCTGCCGGCTACACCTGGGCCATCGCTTGGGCTGCTTTTGGTCTGGCCCTTGGAGGAAACGGGTGGTCACTGTCTTTTAGAG
GCTGCCATATCCCCTTCCAGAAAACAGTTCTCGTTTGTTTACTCGTGGTTCTTGTTCACCCGAGCCTTCTGTTGCCTTTCAGAACTCGCAAACTTTGAGAAGAACGTGAGCCAAGCTATCCACAAGTACAATGCTTACAG
GTGGGACAGTGCAGCATTCTCGGGTAGCATACGTTCTGGGATACCCTGTTTAGTGTGGCAATTAACAGGACTGAGGGCCCAGTGGATATTTGGTCCATCTGCAAGAGCGGGAAAAAGCAAGAATCGAGGCTGGTA
CCTTACTATTTCTTGAAGAATGTGGGCAGTGCGTTATAGGATCAGTTCAGATAAGTTCCCAATTCTGATTGCAGATAGCTGTAGCCTGATACGTATTTTGTCATCATCAAACATGAATTACTAGAAATGCGGTGTAATTT
TAACCGTAATGAAAGACAGCGGATGAAGTCCATACTGTGAAATATATGTGGTTCAAGGAATCACATTTTTTCTTTTGGTTGGTCTAAAGTTGATCCTGTGGGATCCTGATCCTGATCCTTGTTCCTT

Allele 3: Partial insertion

TTGTAAAATAAACTATTGCTGATGTGACATTTCCATTTCTCAAGCAGACTGTAAATTTCTCAGATGTAGGTCATCATGTTATAATTAGGGTTCCATTGCTCATGATCTAGATGAAACACTTCTCCCTTTCTTGTGGTCTCTT
CATTAATTTGGTCCCTACCGCAAAGGCAGGGGGATTAAAAATAAAATATCTTTTGACACTGGTTGGCAGGTGTACTAACACGTGTGCGGGAGGATTCCTGTTCTCGGCATGGTTCACGCCGGGCTCCGGCAGCGCC
CTAGTACGTAAGGGGCTGCACTGGAGTGTGGACTGCCCGGTATCCTTAGAGAAAGTTTTGACAGTGTGACGCCAGTGTCGCCTCTAACCACGTACAAAACGCGCCATGCATTGGGTAGCTATTACAGTGCACATTTT
TGTGCAGGAAGCTACCCCACCAGCGCAAGTATTTCAGAATCAAGATCGCACTCCCGTTTCCCCTTCTAGGGAAAGGATTCCAGATAAACACTGAGAAAACAGATTTAATATCTTAGGTCAAAACTCCAGGTCTTCCCA
TAGGAAGGCCCCTGGGCTGTCATTCTGAGGTGCCTATTTCCCCCGCGCCGTGTCCACGTCCACGAGTCCACGAACCCTCCGAGTTCTCTCTCCTCACCTGCCGAGGGCTCGCCCCTCTAGCCCCGCCCCGCCCCGC
CCAGGACGCGTGACGTCACAACAAGCGCCGTGCGCCCCGCCCCCCATCGGGGGCAACCATTGTTCCGCCGGTCGCGCCGGAGCTGGGTTGCTCCTGCTCCCGTCTCCAAGTCCTGGTACCTCCTTCAAGCTGGGA
GAGGGCTCTAGTCCCTGGTTCTGAACACTCTGGGGTTCTCGGGTGCAGGCCGCCGTGACCGCCGCCGGGATCACTCTCGGCATGGACGAGCTGTACAAGATGAGCAAACGGAAGGCGCCGCAAGAGACTCTCAA
CGGGGGAATCACCGACATGCTCACAGGTTAGCACCGGGCCGGGCCCCGCTGGCTTTCTTCTTTCCTTCCAGCCTCTTCCCCTGCCTTCCTTCTCTCCCACACCGACAGTCCAGTGGGTAGGGTAGGTTCCTTGCAGC
GGGTCGTCTTCCGTGGGGATCTCCCTCCGGCGCCCCTGGCTGGTTGTCAGTCCTGCCGGCTACACCTGGGCCATCGCTTGGGCTGCTTTTGGTCTGGCCCTTGGAGGAAACGGGTGGTCACTGTCTTTTAGAGGCT
GCCATATCCCCTTCCAGAAAACAGTTCTCGTTTGTTTACTCGTGGTTCTTGTTCACCCGAGCCTTCTGTTGCCTTTCAGAACTCGCAAACTTTGAGAAGAACGTGAGCCAAGCTATCCACAAGTACAATGCTTACAGGT
GGGACAGTGCAGCATTCTCGGGTAGCATACGTTCTGGGATACCCTGTTTAGTGTGGCAATTAACAGGACTGAGGGCCCAGTGGATATTTGGTCCATCTGCAAGAGCGGGAAAAAGCAAGAATCGAGGCTGGTACC
TTACTATTTCTTGAAGAATGTGGGCAGTGCGTTATAGGATCAGTTCAGATAAGTTCCCAATTCTGATTGCAGATAGCTGTAGCCTGATACGTATTTTGTCATCATCAAACATGAATTACTAGAAATGCGGTGTAATTTTA
ACCGTAATGAAAGACAGCGGATGAAGTCCATACTGTGAAATATATGTGGTTCAAGGAATCACATTTTTTCTTTTGGTTGGTCTAAAGTTGATCCTGTGGGATCCTGATCCTGATCCTTGTTCCTT
